# Mucosal architecture, microbiota resilience and metabolic disruption reveal holobiont dynamics during bovine respiratory disease

**DOI:** 10.64898/2026.10.02.756151

**Authors:** Sergio Méndez, Güénaelle Derolez, Florence Mompart, Maverick Monie--Ibanes, Thomas Peyret, Marie Corbin, David Bars-Cortina, Fatima-Zohra Sikht, Anne Geffré, Gwendoline Pot, Hortensia Robert, Yves Farizon, Christine Citti, Eric Richard, Laurent Xavier Nouvel, Christine Jansen, Katarzyna Dudek, Christiane Schnee, Eric Baranowski, Gilles Meyer, Mariette Ducatez, Núria Mach

## Abstract

The gut–airway axis is increasingly recognised as an important determinant of respiratory health in livestock. However, the interactions among respiratory infections, microbial communities, mucosal barrier function, and microbial metabolism that underpin this axis remain poorly understood. Here, we combined longitudinal immunological, mucosal, microbiological, and metabolic analyses to characterise host–microbiota interactions during antibiotic treatment and experimental bovine respiratory disease (BRD). We found that the bovine respiratory tract is organised as a hierarchically structured mucosal ecosystem, with compartment-specific mucin expression patterns associated with distinct microbial communities. Within this framework, increasing respiratory pathogen loads were associated with greater clinical severity, oxidative stress, acute inflammation, and targeted alterations in mucosal regulation and microbial composition, while serum amyloid A emerged as the host biomarker most closely reflecting infection intensity. Although infection altered host and microbial states, antimicrobial treatment represented the dominant ecological perturbation. Antibiotic exposure reshaped both respiratory and intestinal microbial communities, reducing diversity, increasing community drift, and driving coordinated ecological transitions across the gut-airway axis that exceeded the effects of infection alone. Importantly, these compositional changes were accompanied by profound reductions in acetate, propionate, butyrate, and valerate, identifying short-chain fatty acids as a major functional consequence of gut microbiota disruption.

Integrative multi-omics analyses revealed that mucosal organisation, microbial diversity, microbial metabolism, and immune activity were embedded within a highly structured gut-airway network. Together, these findings support a holobiont view of BRD in which disease outcomes emerge not only from pathogen-host interactions but also from ecological disturbances that propagate across interconnected mucosal compartments. Notably, antimicrobial treatment affected ecosystem structure and function more than infection itself, highlighting potential trade-offs between pathogen control and microbiota stability. Preserving microbial resilience and its metabolic functions may therefore represent an important component of future strategies to improve respiratory health and disease resilience in cattle.

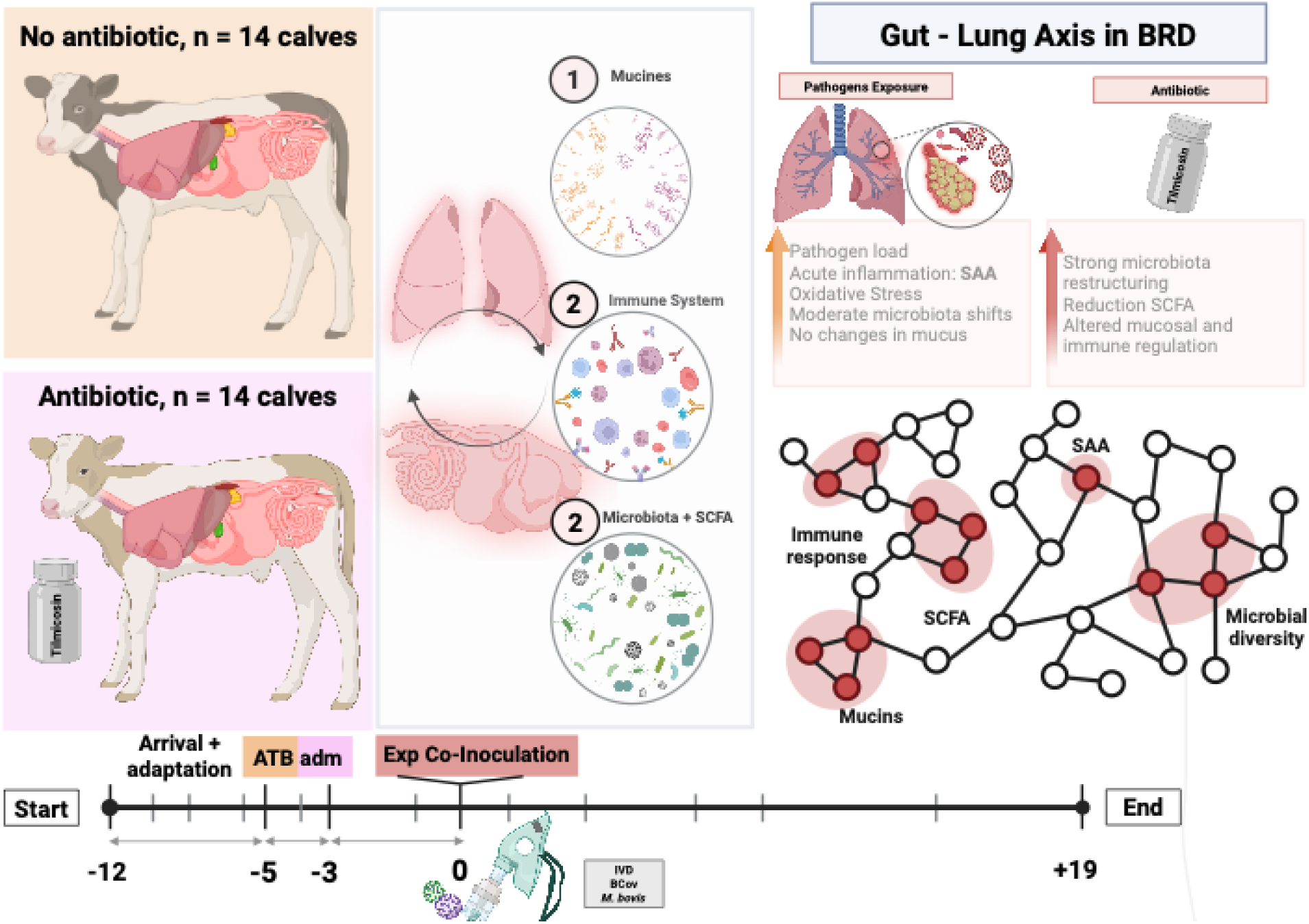

## INTRODUCTION

The concept of the holobiont has profoundly transformed our understanding of animal biology, shifting from a host-centric paradigm toward an integrative view in which the host and its associated microbial communities function as a unified ecological and evolutionary entity [1, 2]. Therefore, health and disease arise not from isolated components but from the dynamic interactions among host tissues, microorganisms, and environmental pressures.

A central yet often underexplored component of the holobiont is the mucosal interface, which constitutes the primary boundary between the host and its surrounding microbial environment [3, 4]. Mucus, largely composed of mucins, a family of highly glycosylated proteins, acts not only as a physical barrier but also as an active biochemical and ecological matrix that governs microbial colonisation, nutrient availability, and host–microbe interactions [5–9]. In humans and experimental animal models, secreted mucins such as MUC5AC and MUC5B, along with membrane-bound mucins (e.g., MUC1, MUC4, MUC16), play critical roles in respiratory pathogen clearance and immune signalling [10, 11]. In the gastrointestinal tract, mucins such as MUC2 form the structural backbone of the mucus layer, shaping microbial colonisation and intestinal homeostasis [12–16]. In cattle, however, mucin biology remains underexplored. A genome-wide analysis of the bovine mucin gene family revealed a diverse repertoire of both membrane-associated and secreted mucins, exhibiting strong regional transcriptional patterns along the gastrointestinal tract, suggesting a highly structured, compartment-specific mucosal barrier system [17]. However, beyond this foundational genomic characterisation, little is known about how mucins are dynamically regulated during infection, how they interact with resident microbiota, or how they respond to antimicrobial perturbations. This gap is especially critical because mucins likely represent a central integrative interface within the holobiont, mediating multidirectional interactions among host tissues, microbial communities, and invading pathogens [3, 4]. Pathogens also exploit and modify this environment to establish infection, while simultaneously triggering host immune responses that reshape mucosal architecture [18–20]. This continuous feedback establishes a multidirectional network in which mucus functions as a central integrative hub linking microbial ecology, pathogen dynamics, and host physiology. Beyond local interactions, mucosal systems are interconnected across distant anatomical sites, forming integrated functional axes such as the gut–lung axis [21, 22]. Growing evidence demonstrates that microbial metabolites, particularly short-chain fatty acids (SCFAs), along with immune mediators and microbial translocation, mediate bidirectional communication between the gastrointestinal and the respiratory tracts [23]. However, how these compartments orchestrate such cross-talk during infection remains poorly understood.

In this context, bovine respiratory disease (BRD) is a good model for studying holobiont dynamics. Despite decades of research, BRD continues to challenge conventional management strategies because of its complex, multifactorial nature, involving viral and bacterial pathogens, host immunity, environmental stressors, and management practices [24–28]. Unlike a single pathogen-driven disease, BRD involves multiple pathogens that interact with resident microbiota and host defences in a dynamic, non-linear manner. Key pathogens include Bovine Respiratory Syncytial Virus (BRSV), Bovine Coronavirus (BCoV), Bovine Parainfluenza-3 Virus (BPIV-3), BHV-1 (Bovine Herpes Virus 1), and BVDV (Bovine Viral Diarrhoea Virus) [29, 30]. Bacterial contributors include *Pasteurella multocida, Histophilus somni,* and *Mannheimia haemolytica*, as well as *Mycoplasma bovis* [25]. Together, they contribute to respiratory epithelial damage, immune dysregulation, and increased susceptibility to secondary infections [28, 31, 32]. Furthermore, BRD affects both the respiratory and gastrointestinal compartments [33, 34], either directly or indirectly, thereby providing a unique opportunity to investigate multi-compartment interactions within the holobiont framework.

A particularly relevant dimension of BRD is its intersection with antibiotic use, a major perturbation to microbial ecosystems worldwide [35, 36]. Antibiotics, while essential for disease control, are among the most significant ecological disturbances imposed on host-associated microbiota [37–39]. Their effects extend far beyond pathogen suppression, often reducing microbial diversity, altering metabolic outputs, and disrupting ecological stability across multiple body compartments [40].

Despite growing recognition of the holobiont, most studies of infectious diseases continue to focus on individual components, either pathogens, microbiota, or host responses, without addressing their integrated and dynamic interplay [25, 30, 41–53]. Addressing these questions requires longitudinal, multi-scale approaches that capture the coordinated responses of host physiology, microbial ecosystems, and mucosal architecture.

In this study, we leverage the BRD model to investigate the holobiont as a multi-compartment, multi-layered system that integrates respiratory and intestinal microbiota, mucin expression, host immune and metabolic responses, and pathogen dynamics. By combining a controlled factorial design that independently manipulates antibiotic exposure and infection with longitudinal sampling across multiple biological compartments, we aim to disentangle the relative contributions of these perturbations and to characterise their combined effects on the holobiont.

## RESULTS

### Study design and integrative framework

To investigate host–microbe–mucus interactions during bovine respiratory disease, we implemented a longitudinal, multi-compartment sampling strategy across respiratory and intestinal sites. Animals were allocated to a study design in which antibiotic exposure and infection status were controlled. Specifically, half of the animals received the macrolide antibiotic tilmicosin from day −5 to day −3, while the remaining animals received no antibiotic treatment. On day 0, within each of these two groups, only half of the animals were experimentally co-inoculated with bovine coronavirus (BCoV), influenza D virus (IDV), and *M. bovis*. This design resulted in four groups: antibiotic + inoculation, antibiotic + no inoculation, no antibiotic + inoculation, no antibiotic + no inoculation, enabling us to disentangle the individual and combined effects of inoculation and antibiotic treatment. Mucin expression and microbiota composition in both nasal and faecal samples, together with host immune and metabolic responses, were profiled over time to monitor the trajectory of the holobiont and capture coordinated changes across these interconnected biological systems (Figure 1A–B; Additional File 1).

**Figure 1.**
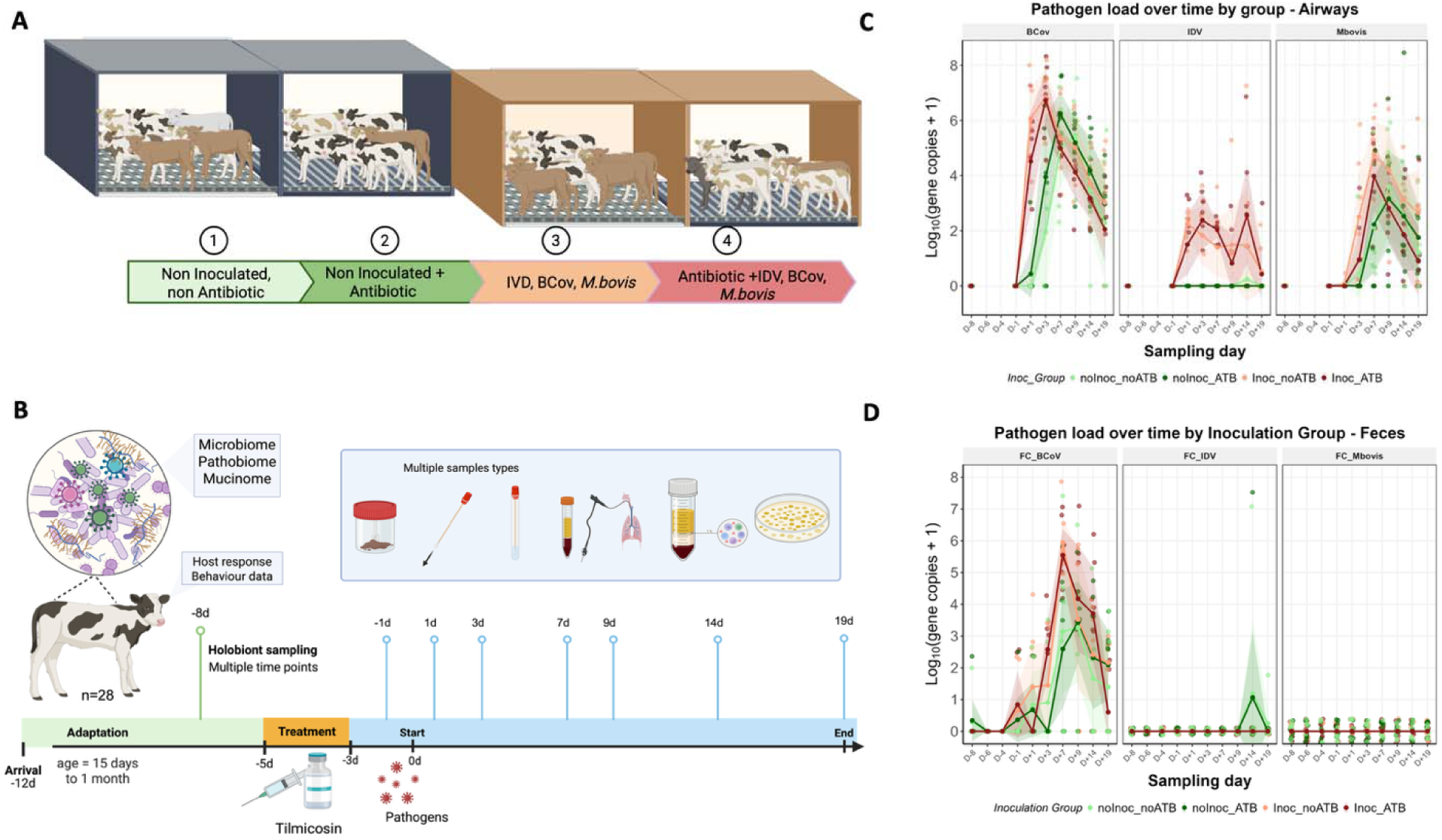
Experimental design, sampling strategy, pathogen dynamics, and clinical outcomes. **(A)** Schematic representation of the experimental setup and group allocation. Calves (n = 28) were distributed int four groups: non-inoculated without antibiotics, non-inoculated treated with tilmicosin, inoculated with IDV, BCoV, and *M. bovis*, and inoculated animals receiving antibiotic treatment; **(B)** Overview of sampling strategy and stud timeline. Following an adaptation period, calves were treated with antibiotics (days −5 to −3) and experimentally inoculated at day 0. Longitudinal sampling across multiple compartments (nasal swabs, faeces, blood) was performed from day −8 to day 19; **(C-D)** Pathogen load dynamics in the airways and faecal samples, respectively, over time. Log -transformed gene copies (+1) of BCoV, IDV, and *M. bovis* are shown across sampling days for each experimental group. Lines represent group-level trends, and points represent individual animals.

### Experimental validation and pathogen loads

Before inoculation, all calves tested negative for the experimental pathogens *M. bovis*, BCoV, IDV, and other major respiratory pathogens, including *Mannheimia haemolytica*, *Pasteurella multocida*, *Histophilus somni*, bovine respiratory syncytial virus (BRSV), and bovine parainfluenza virus type 3 (BPIV-3). Pathogen loads in the nasal cavity increased following challenge with IDV, BCoV and *M. bovis*, supporting exposure and replication of the inoculated pathogens (Additional File 2).

Specifically, qPCR rapidly detected IDV and BCoV in the nasal cavity of inoculated animals from day 1 post-infection, whereas *M. bovis* showed delayed kinetics, becoming detectable from day 3 and declining thereafter (Figure 1C, *p* < 0.05). Serological evidence further supported establishment of *M. bovis* infection, as several challenged animals seroconverted during the study. Although all animals were seronegative before inoculation, we detected *M. bovis*-specific antibodies in a subset of animals at Day +19.

However, pathogen detection was not confined to inoculated animals. Nasal BCoV was detected in non-inoculated calves from day 3, and *M. bovis* from day 7, indicating progressive within-cohort transmission despite biosecurity measures. Culture, PCR, and whole-genome sequencing of *M. bovis* isolates recovered from the nasal cavity of non-inoculated animals confirmed that they originated from the inoculated RM16 strain. Detected variants were largely attributable to low-frequency polymorphisms already present in the inoculum, with no evidence of unrelated *M. bovis* strains, thereby confirming transmission within the experimental population.

Faecal pathogen analyses revealed delayed dynamics relative to those observed in nasal samples. BCoV and IDV were detected in faecal samples throughout the study period (Figure 1D), while *M. bovis* was not detected. As in the respiratory tract, BCoV and IDV were found in both challenged and non-challenged animals.

### Heterogeneous infection dynamics motivate an individual-level, pathogen load–based framework

Because natural transmission occurred within the cohort, inoculation profiles overlapped between experimentally inoculated and non-inoculated animals, limiting the relevance of group-based classifications. We therefore performed subsequent analyses at the individual level, using pathogen load as a continuous variable. We generated composite pathogen load indices from quantified respiratory and enteric inoculated pathogens to capture the overall infectious pressure each animal experienced over time. Consequently, pathogen load represented biological exposure better than treatment group assignment. This approach enabled us to investigate host, microbial, and mucosal responses across a gradient of pathogen abundance, providing a biologically relevant framework for studying host–microbe–mucus interactions during BRD.

#### Pathogen abundance drives disease severity and acute inflammation, but has limited effects on adaptive immune responses

Increasing pathogen load was strongly associated with disease severity and acute inflammatory responses, whereas adaptive immune and haematological changes were largely structured by time post-challenge.

Clinical scores increased with pathogen abundance, particularly between D+3 and D+13 (Figure 2AB). While blood lactate concentrations varied over time, they were unrelated to pathogen counts (β = −0.008, *P* = 0.483). In contrast, total antioxidant status (TAS) declined after infection and was negatively associated with pathogen abundance (β = −0.0026, *P* = 0.0038; Figure 2C).

**Figure 2.**
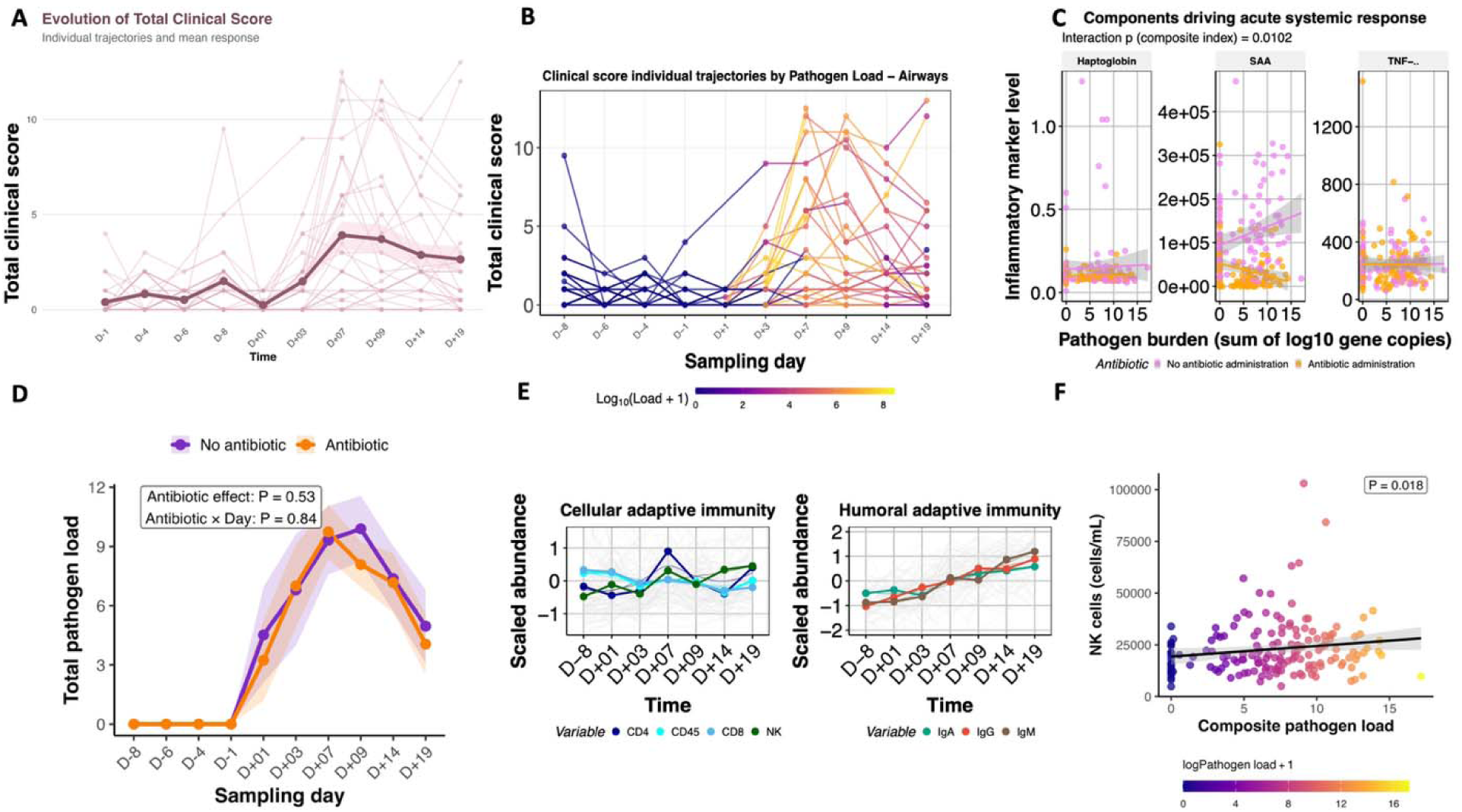
Host physiological, inflammatory and immune responses to respiratory pathogen burden. **(A)** Individual clinical score trajectories throughout the infection challenge; **(B)** individual clinical score trajectories coloured according to pathogen burden. Each point represents an individual; **(C)** Acute inflammatory responses as a function of pathogen concentration; **(D)** Pathogen abundance across time in treated and untreated individuals; **(E)** Trajectories of cellular (CD4, CD8, CD45 and NK cells) and humoral (IgA, IgG and IgM) adaptive immun responses over time; **(F)** Relationship between composite pathogen load and NK cell abundance. NK cell counts increased with pathogen burden.

Acute-phase proteins showed the strongest relationships with pathogen abundance. Haptoglobin concentrations were influenced by the interaction between pathogen abundance and antibiotic treatment (slope = −0.01, *p* = 6.5 × 10□□; Figure 2C). A stronger treatment-dependent pattern emerged for serum amyloid A (SAA), whose relationship with pathogen abundance was significantly modified by antibiotic exposure (estimate = −0.068 ± 0.026, *p* = 0.010). In untreated calves, SAA increased with pathogen concentration (estimate = 0.037 ± 0.018, *p* = 0.043), peaking between D+03 and D+09, whereas this association was reversed in treated animals. Notably, SAA was not associated with any individual pathogen (*M. bovis*, *p* = 0.18; BCoV, *p* = 0.07; IDV, *p* = 0.095), suggesting that it reflected the cumulative inflammatory consequences of polymicrobial infection rather than the effect of a single agent. In contrast, TNF-α was unaffected by pathogen load (*p* = 0.92) or by the pathogen abundance × antibiotic interaction (*p* = 0.67; Figure 2C).

Importantly, these inflammatory patterns could not be attributed to differences in pathogen load between antibiotic treatment groups, as neither antibiotic treatment (F□,□□.□ = 0.40, *p* = 0.53) nor its interaction with sampling day (F□,□□□.□ = 0.55, *p* = 0.84; Figure 2D) affected total pathogen load. This finding is consistent with the antimicrobial susceptibility profile of the inoculated *M. bovis* isolate, which was resistant to tilmicosin at concentrations exceeding 250 µg/mL.

In contrast, adaptive immune responses were only weakly associated with pathogen abundance. Significant pathogen load × time interactions were observed for CD45 cells (F□,□□□.□ = 2.76, *p* = 0.021) and CD4 T cells (F□,□□□.□ = 2.31, *p* = 0.048; Figure 2E). However, pathogen counts alone did not affect cell abundance, except for NK cells (F□,□□□.□ = 5.70, *p* = 0.018; Figure 2F). Likewise, humoral responses were driven primarily by time, with strong temporal effects on IgG (F□,□□□□.□ = 9.78, *p* < 0.001) and IgM (F□,□□□.□ = 11.82, *p* < 0.001), and a marginal effect on IgA (F□,□□.□ = 2.01, *p* = 0.081; Figure 2E). Although pathogen concentration significantly affected IgG (F□,□□□.□ = 6.16, *p* = 0.014), direct associations were weak (β = 0.64-0.73, *p* = 0.094-0.11), and only modest positive relationships were detected for IgA (β = 0.21-0.33, *p* = 0.049 and 0.0065) and IgM (β = 0.41-0.57, *p* = 0.023 and 0.035).

Haematological responses followed a similar pattern (Figure 3A). Leukocyte counts declined with increasing pathogen counts in treated calves (β = −0.11, *p* = 0.004), whereas this relationship disappeared in untreated animals (β = −0.05, *p* = 0.29). Neutrophils showed the strongest association, decreasing with pathogen loads (β = −116.8, *p* = 4.8 × 10□□), but again only in treated calves (untreated: β = −29.3, *p* = 0.43). Correlations between conventional haematology and flow cytometry indicated only partial overlap, with lymphocyte counts moderately correlated with CD45 (*r* = 0.57), CD8 (*r* = 0.41), and CD4 cells (*r* = 0.27), whereas neutrophils and monocytes showed weak correlations with cytometric markers (*r* ≤ 0.24; Figure 3B).

**Figure 3.**
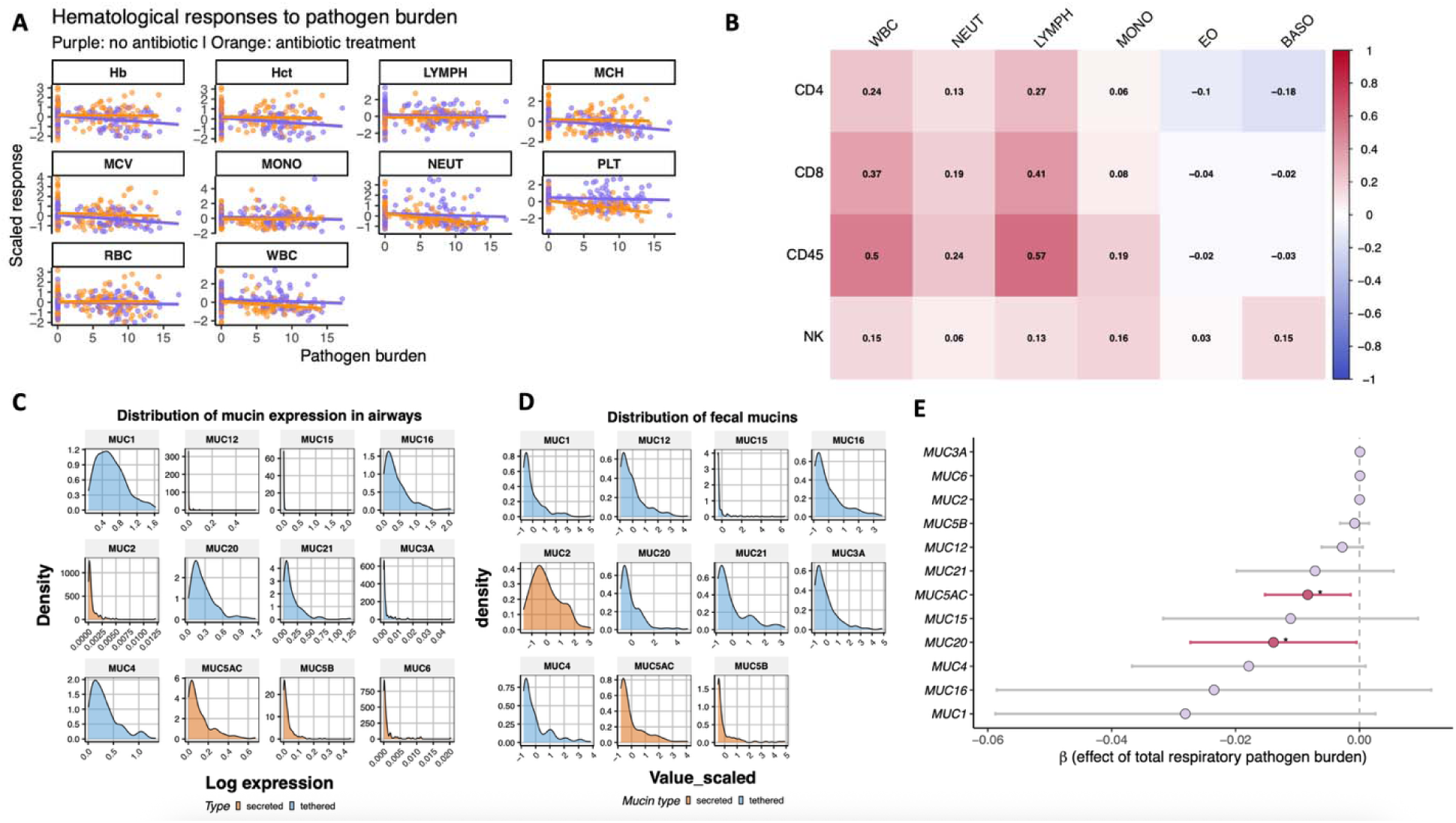
Respiratory pathogen load is associated with selective components of the immune response and mucin expression dynamics. **(A)** Linear regressions between pathogen load and selected immune an haematological variables, shown separately for antibiotic-treated (orange) and untreated (purple) calves; **(B)** Heatmap of Pearson correlation coefficients for immune cells detected by cytometry or via haematology; **(C-D)** Distribution of airway and faecal mucin expression levels, shown as density plots for individual mucins. Secrete mucins are highlighted in orange, and membrane-associated (tethered) mucins in blue. Expression values are log-transformed; **(E)** Association between total respiratory pathogen burden and mucin expression. Forest plot showing β estimates and 95% confidence intervals from linear mixed-effects models evaluating the relationship between total respiratory pathogen burden and airway mucin expression. Negative β values indicate reduced mucin expressio with increasing respiratory pathogen burden, whereas positive values indicate increased expression.

### Distinct mucus architectures govern airway and intestinal responses to infection and antibiotic perturbation

Because mucus constitutes the primary interface between host tissues, pathogens, and microbial communities, we first sought to characterise the organisation and dynamics of mucins across the respiratory and intestinal compartments. To our knowledge, this represents the first comprehensive analysis of airway mucin expression in calves during experimental respiratory infection and antibiotic exposure.

In the respiratory tract, membrane-associated mucins dominated, particularly MUC1, MUC4, MUC16, MUC20, and MUC21, whereas secreted gel-forming mucins contributed comparatively little to overall expression (Figure 3C; Additional File 3). Among secreted mucins, MUC5AC displayed substantially higher expression than MUC2, MUC5B and MUC6. In contrast, the intestinal mucus layer exhibited a fundamentally different architecture, characterised by overwhelming dominance of MUC2, which was expressed at substantially higher levels than all other mucins (Figure 3D). Within the tethered mucins, MUC1, MUC4, MUC12, MUC16, and MUC21 displayed the highest expression and broadest distribution across samples, while MUC15, MUC20, and MUC3A were generally less abundant.

Notably, the respiratory mucosal barrier proved remarkably dynamic and responsive to perturbations. Respiratory pathogen burden was associated with a broad downregulation of airway mucin expression (Figure 3E). Increasing pathogen load significantly reduced the expression of MUC20 (β = −0.0139, *p* = 0.042) and the secreted mucin MUC5AC (β = −0.0083, *p* = 0.018), while similar negative trends were observed for MUC1 (*p* = 0.072), MUC4 (*p* = 0.063), and MUC12 (*p* = 0.101). Notably, pathogen-specific analyses identified no significant effects of *M. bovis*, IDV, or BCoV individually after multiple-testing correction (all FDR-adjusted *p* = 0.996). In contrast, antibiotic treatment markedly remodelled the airway epithelial surface, as shown by a significant time × antibiotic interaction (Figure 4AB). Antimicrobial exposure increased the expression of several membrane-associated mucins, including MUC1 (β = 0.515, *p* = 0.0062), MUC4 (β = 0.571, *p* = 0.0061), MUC16 (β = 0.560, *p* = 0.0017), MUC20 (β = 0.398, *p* = 0.021), and MUC21 (β = 0.358, *p* = 0.031). This response was particularly pronounced at D−1, immediately following antibiotic administration, when all five mucins were simultaneously upregulated (Figure 4A).

**Figure 4.**
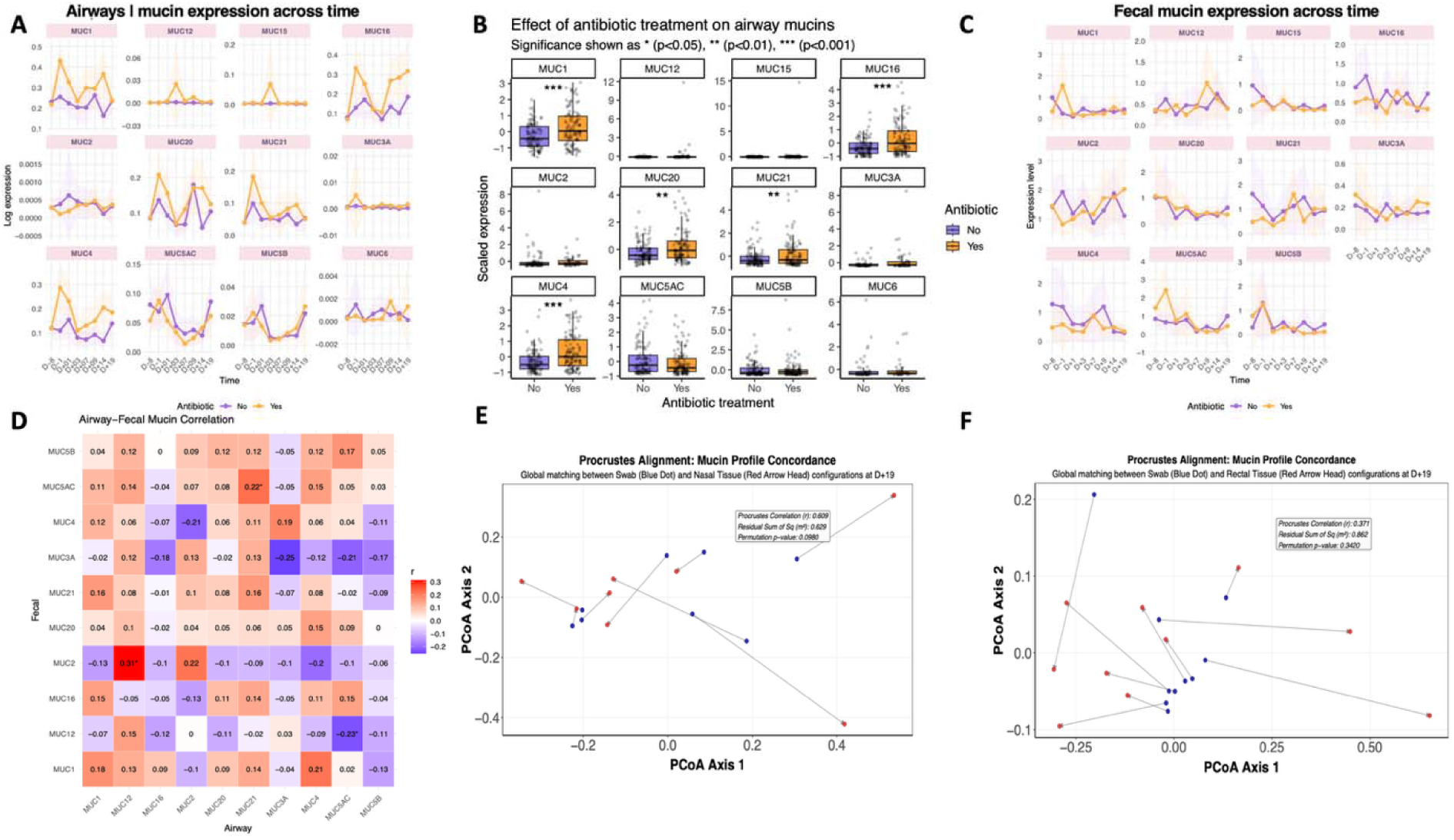
Mucin expression profiles in airways and faeces and their modulation by antibiotics and pathogen burden. **(A)** Longitudinal dynamics of airway mucin expression across time. Lines represent group-level trends (mean ± confidence interval), stratified by antibiotic treatment (orange: treated; purple: untreated), with individual trajectories shown in the background; **(B)** Effect of antibiotic treatment on airway mucin expression. Boxplots represent scaled expression levels in antibiotic-treated versus untreated groups; **(C)** Longitudinal analysis of rectal mucin expression across time according to antibiotic treatment (orange: treated; purple: untreated). Shaded areas represent confidence intervals around the mean expression; **(D)** Heatmap showing Pearson correlation coefficients between airway and faecal mucin gene expression profiles. Positive correlations are shown in red and negative correlations in blue, with correlation coefficients displayed within each cell. Significant associations are indicated by asterisks (*P* < 0.05); **(E)** Procrustes alignment between nasal swab (blue) and nasal turbinate tissue (red) muci profiles, and **(F)** between rectal swab and rectal tissue. Points represent individual samples projected in PCA space, and connecting lines indicate the displacement required to align corresponding samples between datasets. The inset reports the Procrustes correlation and alignment significance.

In contrast, the intestinal mucus barrier remained comparatively stable. Increased faecal pathogen load was associated with reduced expression of MUC16 (β = −0.088, *p* = 0.026), MUC5B (β = −0.129, *p* = 9.5 × 10□□), and MUC5AC (β = −0.103, *p* = 0.038), while greater respiratory pathogen count was linked to lower intestinal expression of MUC20 (β = −0.0068, *p* = 0.020), MUC1 (β = −0.0064, *p* = 0.029), and MUC5AC (β = −0.0099, *p* = 0.006). Antibiotic treatment exerted only modest effects, reducing MUC16 (β = −0.055, *p* = 0.039), MUC15 (β = −0.062, *p* = 0.049), and MUC5B (β = −0.054, *p* = 0.034). Although temporal variation was detected for several accessory mucins, including MUC5AC (*p* < 0.001), MUC5B (*p* = 0.0016), MUC15 (*p* = 0.029), and MUC20 (*p* = 0.034), the dominant structural mucin MUC2 remained unaffected by time, pathogen loads, or antibiotic treatment (all *p* > 0.26; Figure 4C).

Despite these coordinated responses to perturbations, cross-compartment mucin relationships were surprisingly limited. Correlation and regression analyses revealed only weak associations between airway and intestinal mucins (Figure 4D). The strongest positive association wa observed between intestinal MUC2 and airway MUC12 (*r* = 0.31) and MUC2 (*r* = 0.22), while weaker correlations were observed for airway MUC5AC (*r* = −0.23) and rectal MUC12. NMDS analyses similarly failed to reveal coherent clustering between respiratory and intestinal mucin profiles.

Lastly, we evaluated whether minimally invasive sampling could reliably capture tissue-level mucin expression patterns. Nasal swab profiles were significantly correlated with those of nasal tissue (Procrustes *r* = 0.609, *p* = 0.009; Figure 4E), particularly for the membrane-associated mucins MUC1, MUC15, MUC20, and MUC21. In contrast, rectal swab profiles did not accurately reflect intestinal tissue patterns (Procrustes *r* = 0.371, *p* = 0.342; Figure 4F). Among individual mucins, rectal MUC16 showed the strongest correlation between swab and tissu measurements (*r* = 0.64, *p* = 0.0004), followed by MUC5B (*r* = 0.41, *p* = 0.036) and MUC2 (*r* = 0.37, *p* = 0.01).

### Antibiotics override pathogen effects in structuring airway and gut microbiota

#### Microbiota composition across the airway and faecal compartments

Sequencing of nasal and faecal samples yielded mean read counts of 36,211.33 ± 1,789.04 and 75,929.23 ± 2,691.96 reads per sample, respectively, after quality filtering. We detected 9,765 ASVs in the airway dataset (Additional File 4) and 3,039 in the faecal dataset (Additional File 5), with mean richness of 218.63 ± 8.22 ASVs per sample in the airway and 159.10 ± 3.77 ASVs per sample in the faecal dataset.

Distinct microbial communities were observed in the airway and faecal compartments. At the phylum level, the airway microbiota was dominated by Proteobacteria (43.2% ± 1.41), followed by Bacillota (38.6% ± 1.38), Bacteroidota (7.61% ± 0.573), Actinomycetota (6.58% ± 0.396), and Fusobacteriota (3.27% ± 0.517). In contrast, the faecal microbiota was primarily composed of Bacillota (55.5% ± 0.849) and Bacteroidota (31.6% ± 0.835), with lower contributions from Actinomycetota (7.73% ± 0.356) and Pseudomonadota (3.76% ± 0.265).

Taxonomic richness further reflected this compartmentalisation. The airway microbiota comprised 371 families and 832 genera and was predominantly driven by a limited number of highly abundant families, including *Mycoplasmataceae* (31.6 % ± 1.44), *Moraxellaceae* (27.1% ±1.42), and *Pasteurellaceae* (6.29% ± 0.575), alongside contributions from *Lachnospiraceae* (12.2% ± 0.486) and *Ruminococcaceae* (10.7% ± 0.442; Figure 5A, Additional File 6). At the genus level, dominant taxa included *Mycoplasma* (31.5% ± 1.43), *Faucicola* (14.3% ± 1.22), *Moraxella* (8.95% ± 0.837), *Caviibacter* (3.13% ± 0.506), and *Mannheimia* (2.88% ± 0.353), together with lower-abundance but recurrent genera such as *Acinetobacter, Pasteurella, Streptococcus*, and *Pseudomonas* (Figure 5B; Additional file 6).

**Figure 5.**
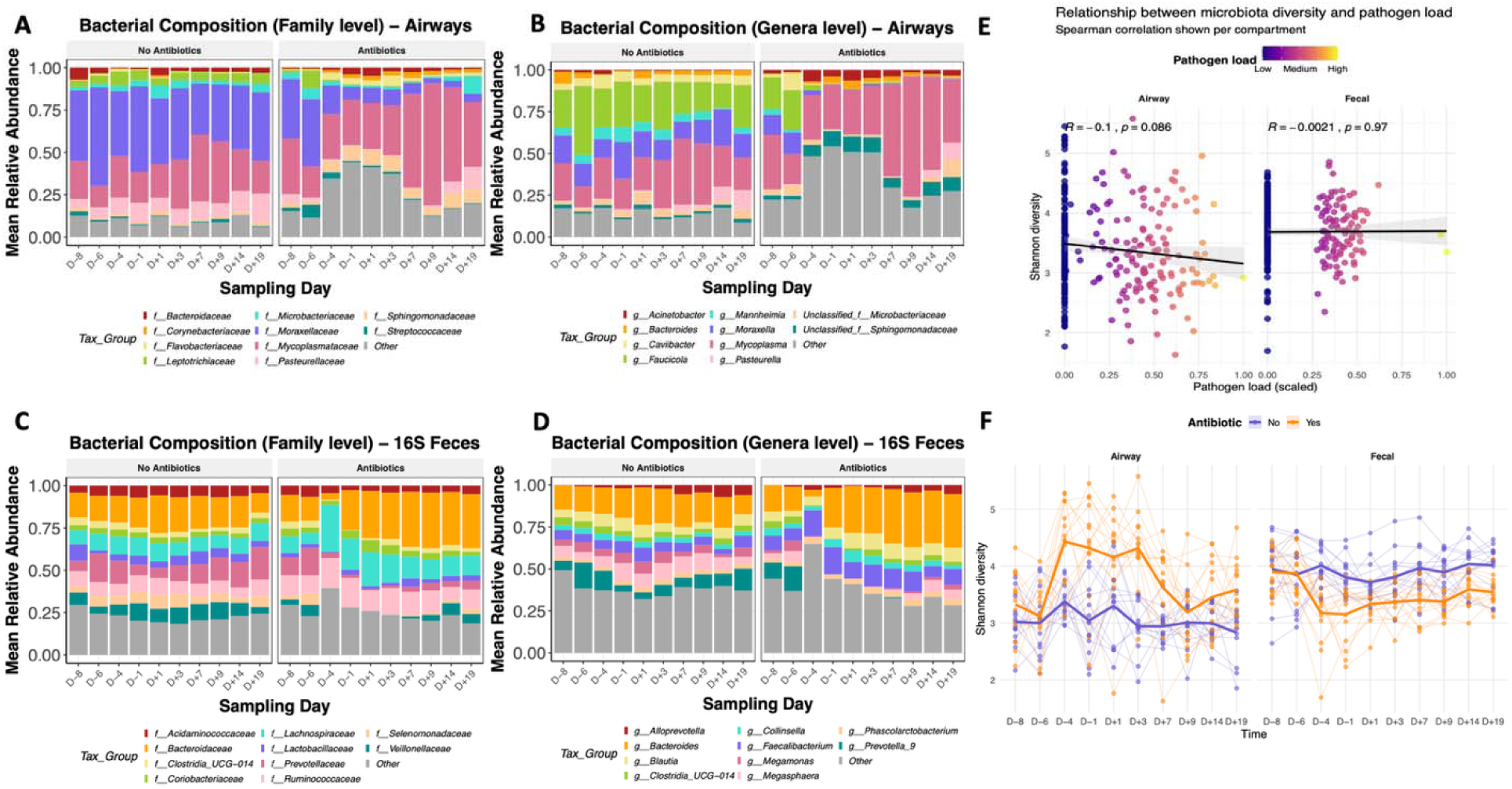
Antibiotic treatment alters airway microbiota composition, diversity and biomass, whereas the faecal microbiota remains comparatively stable. **(A)** Mean relative abundance of the dominant bacterial families in the airway microbiota across sampling days in calves receiving no antibiotics (left) or antibiotics (right). Stacked bars represent mean community composition at each time point; **(B)** Mean relative abundance of the dominant bacterial genera in the airway microbiota across sampling days by antibiotic treatment. Antibiotic administration was associated with marked shifts in taxonomic composition over time; (**C)** Mean relative abundance of the dominant bacterial families in the faecal microbiota across sampling days in calves receiving no antibiotics (left) or antibiotics (right). Stacked bars represent the mean community composition at each time point; **(D)** Mean relative abundance of dominant bacterial genera in the faecal microbiota across sampling days by antibiotic treatment. Antibiotic administration was associated with marked shifts in taxonomic composition over time; **(E)** Relationship between scaled pathogen load and Shannon diversity in the airway and faecal microbiota. Each point represents an individual sample and is coloured by pathogen burden; **(F)** Temporal dynamics of Shannon diversity in airway and faecal microbial communities by antibiotic treatment. Thin lines represent individual calves and thick lines represent group means.

In contrast, the faecal microbiota exhibited lower taxonomic richness, with 160 families and 303 genera, and was dominated by gut-associated taxa, particularly *Bacteroidaceae* (20.8% ± 0.815; Figure 5C). Dominant genera included *Bacteroides* (2.44% ± 0.367), along with typical commensal and fermentative taxa such as *Faecalibacterium, Blautia, Dorea, Streptococcus*, and *Escherichia-Shigella,* reflecting a structured and functionally specialised gut ecosystem (Figure 5D; Additional File 7).

Core microbiota analysis at the genus level further emphasised differences in ecological composition between compartments. In the airway, only a limited set of genera consistently occurred across samples, including *Flavobacterium, Mycoplasma, Corynebacterium, Moraxella, Psychrobacter*, and *Bifidobacterium*, indicating a relatively constrained yet variable core community (Additional File 8). Given *Mycoplasma*’s prominence as a core airway genus, we further characterised cultured isolates recovered from nasal swabs and BAL at the species level using 23S rRNA gene sequencing. This approach showed that *M. bovis* was never detected as a single-species coloniser but consistently co-occurred with other *Mycoplasma* species. The most common co-occurring species were *M. dispar*, detected in 45% of *M. bovis*-positive samples, and *M. bovoculi*, detected in 35%. However, no specific *Mycoplasma* species showed a systematic association with *M. bovis*.

In contrast, the faecal microbiota showed a more robust and conserved core of genera such as *Bacteroides, Faecalibacterium, Streptococcus, Dorea, Blautia*, and *Escherichia-Shigella*, consistent with the greater stability typically observed in the gut microbiota (Additional File 9).

#### Antibiotic treatment, rather than pathogen abundance, drives microbiota restructuring across the gut–airway axis

To assess how microbial communities responded to respiratory infection and antibiotic exposure, we analysed microbial diversity, biomass, community composition, and temporal stability using absolute abundance data, avoiding the compositional constraints of relative abundance analyses. In the airway microbiota, Shannon diversity was not associated with pathogen counts (F = 0.06, *p* = 0.81), although a weak negative trend was observed (Figure 5E). In contrast, Shannon diversity was strongly influenced by antibiotic exposure (F = 40.93, *p* = 9.45 × 10□□) and time (F = 7.60, *p* = 9.72 × 10 ¹□□), with a significant antibiotic × time interaction (F = 4.25, *p* = 4.18 × 10□□; Figure 5F; Additional File 10). The changes in airway diversity was accompanied by a marked decline in bacterial biomass (F□,□□.□ = 27.95, *p* = 2.22 × 10□□), corresponding to an average decrease of 0.51 log bacterial gene copies per gram in antibiotic-treated calves, whereas pathogen burden had no effect on biomass (F□,□□.□ = 0.92, *p* = 0.339; Figure 6A).

**Figure 6.**
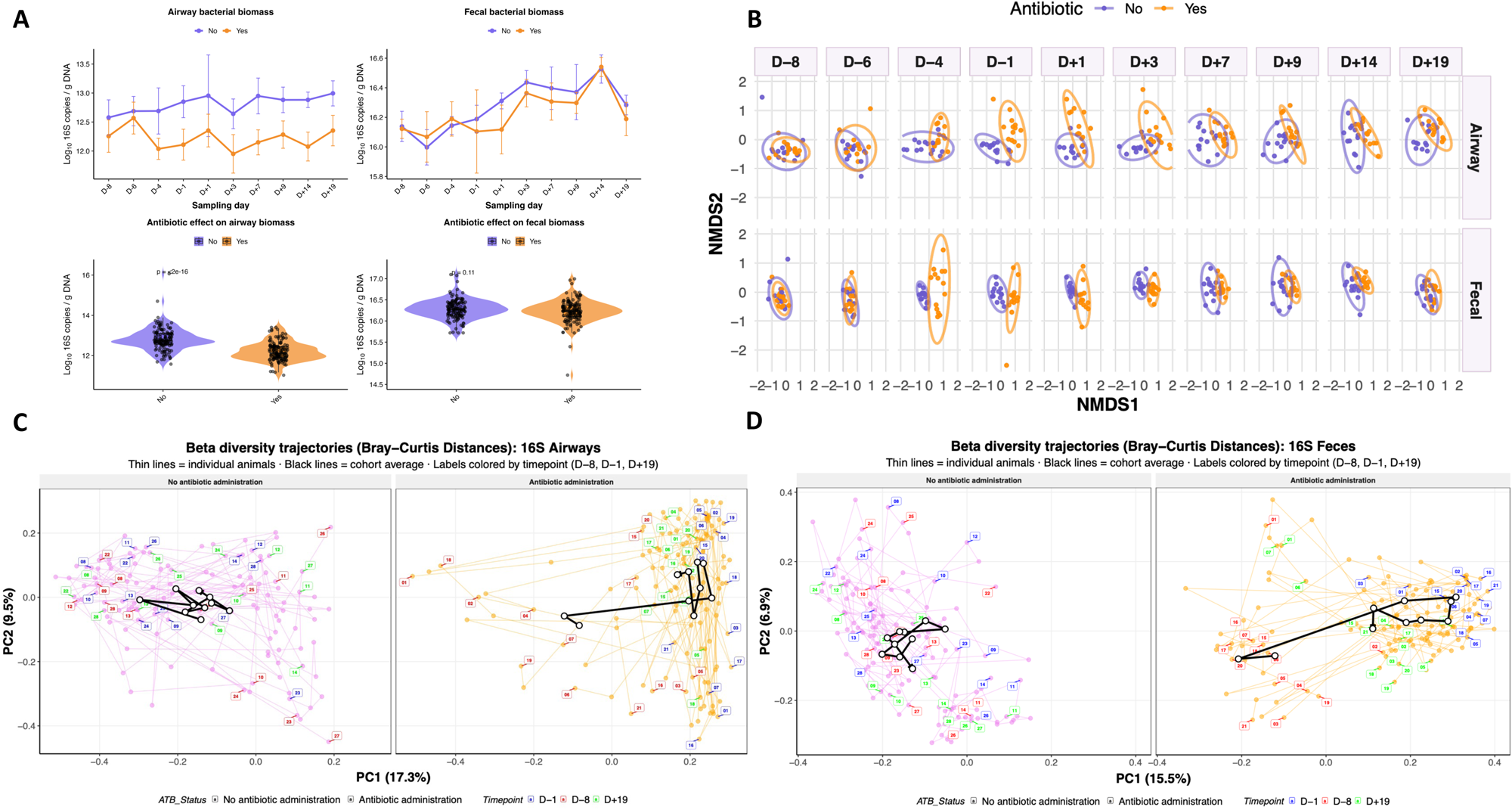
Beta-diversity dynamics and coordinated airway–faecal microbiota responses over time. **(A)** Temporal variation in total microbial biomass in the airway and faecal compartments according to antibiotic treatment. Lines represent group means and shaded areas represent confidence intervals; **(B)** NMDS ordination showing temporal changes in airway and faecal microbiota composition across the study period. Samples are grouped by sampling time, from D−8 to D+19; **(C)** Faecal microbiota trajectories and **(D)** airway microbiota trajectories based on Principal Coordinates Analysis (PCoA) of Bray–Curtis dissimilarities. Each trajectory represents an individual animal, with coloured labels denoting the sampling time point (D−8, D−1 and D+19). Thin lines connect repeated samples from the same animal, while black lines represent the average trajectory of each treatment group.

A similar pattern was observed in the faecal microbiota. Shannon diversity was significantly affected by antibiotic treatment (F = 19.81, *p* = 1.45 × 10□□), time (F = 5.77, *p* = 3.15 × 10□□), and their interaction (F = 3.87, *p* = 1.39 × 10□□; Figure 5F), whereas pathogen load remained unrelated to diversity (F = 0.02, *p* = 0.89; Figure 5E; Additional File 11). However, unlike the respiratory ecosystem, faecal bacterial biomass remained remarkably stable and was unaffected by either antibiotic treatment or pathogen abundance, varying only through time (Figure 6A).

Community composition analyses further highlighted antibiotics’ dominant role in shaping microbiota dynamics. NMDS ordination revealed a strong separation between respiratory and intestinal communities, with compartment representing the principal source of variation (PERMANOVA: R² = 0.202, *p* = 0.001; Figure 6B). Within each ecosystem, antibiotic treatment consistently explained more community variation than pathogen burden. In the airways, antibiotics accounted for 8.2% of compositional variation (R² = 0.082, *p* = 0.001), whereas pathogen abundance explained only 1.3% (R² = 0.013, *p* = 0.001). Likewise, in the faecal microbiota, antibiotic exposure explained 7.7% of variation (R² = 0.077, *p* = 0.001), compared with only 2.6% for pathogen load (R² = 0.026, *p* = 0.001).

Longitudinal trajectory analyses reinforced these findings. Untreated animals followed relatively constrained paths through ordination space, whereas antibiotic-treated animals deviated more from their baseline microbiota and showed substantially greater inter-individual variability (Figure 6C–D). The centroids of treated communities progressively diverged from their pre-treatment position in both compartments, indicating sustained ecological restructuring following antibiotic administration.

Consistent with these compositional shifts, temporal stability analyses showed progressive microbiota divergence throughout the experiment (*p* < 2.2 × 10 ¹□□), amplified by antibiotic treatment in both ecosystems (airway: *p* = 0.008; faecal: *p* = 0.0004). This reduction in stability was accompanied by a decline in community resistance, indicating reduced microbial resilience following antibiotic exposure (Figure 7A). Destabilisation occurred earlier and more abruptly in the faecal microbiota, whereas the airway microbiota showed a more moderate but prolonged perturbation before gradually returning toward baseline.

**Figure 7.**
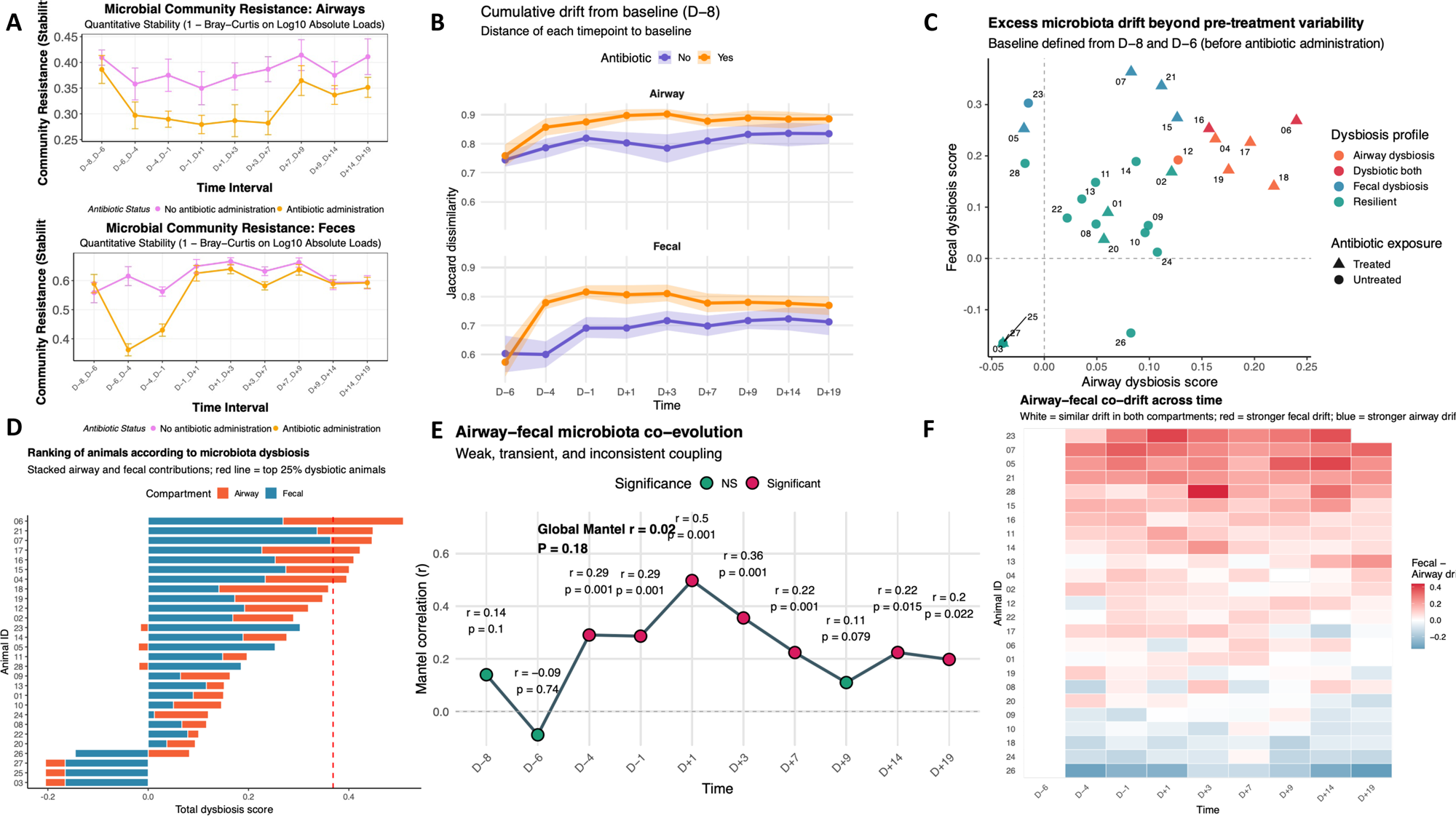
Temporal dynamics, resilience and coordination of airway and faecal microbiota following antibiotic administration. **(A)** Community resistance. Quantitative community resistance was calculated as 1 Bray–Curtis dissimilarity using log-transformed absolute microbial loads between consecutive sampling intervals. Values closer to 1 indicate greater stability, whereas lower values indicate stronger community perturbation. Resistance was assessed separately for airway (upper panel) and faecal (lower panel) microbiota. Points represent group means and error bars indicate standard errors. Purple lines correspond to untreated animals and orange lines to antibiotic-treated animals; **(B)** Cumulative drift from baseline. Temporal divergence of microbial communities from the pre-treatment baseline sample collected at D−8, measured using Jaccard dissimilarity. Higher values indicate a greater departure from the initial microbiota composition. Drift was evaluated separately for airway (upper panel) and faecal (lower panel) microbiota. Lines represent mean values, and shaded ribbons represent confidence intervals. Purple and orange lines indicate untreated and antibiotic-treated animals, respectively; **(C)** Individual dysbiosis profiles. Scatterplot showing animal-specific dysbiosis scores in the airway (x-axis) and faecal (y-axis) microbiota. Dysbiosis was defined as excess post-treatment microbiota drift relative to pre-treatment variability measured between D−8 and D−6. Each point represents one animal, and symbols indicate antibiotic exposure (circles: untreated; triangles: treated). Colours indicate dysbiosis categories: resilient, airway dysbiosis, faecal dysbiosis, or dysbiosis in both compartments; **(D)** Ranking of animals according to dysbiosis. Animals were ranked by total dysbiosis score. Stacked bars show the respective contributions of airway (orange) and faecal (blue) microbiota to the overall dysbiosis burden. The red dashed line indicates the threshold corresponding to the top 25% most dysbiotic animals; **(E)** Airway–faecal microbiota co-variation assessed by Mantel correlation across time points. Correlation strength (r) and significance are shown for each time point; **(F)** Heatmap showing the difference between faecal and airway microbiota drift from baseline (Δ drift = faecal drift − airway drift) for each animal across time. White values indicate similar drift in both compartments, red values indicate greater drift in the faecal microbiota, and blue values indicate greater drift in the airway microbiota. Animals are ordered by their average Δ drift over the study. The predominance of values near zero indicates that both microbial ecosystems generally drifted in parallel over time.

#### Inter-individual susceptibility to dysbiosis reveals resilient and vulnerable holobionts

Dysbiosis, quantified as the microbiota drift observed after antibiotic administration relative to pre-treatment variability (D−8 to D−6), was strongly associated with antibiotic exposure (χ² = 16.25, df = 3, *P* = 0.001), confirming that tilmicosin was the principal driver of microbiota destabilisation across both respiratory and intestinal ecosystems (Figure 7CD).

Despite this overall treatment effect, responses varied substantially among calves (Figure 7D). The most dysbiotic animals (06 and 16) exhibited pronounced microbiota drift in both the respiratory and intestinal compartments, whereas resilient animals (03, 25, and 27) maintained microbial communities close to their pre-treatment state (Figure 7D). Notably, calf 03 displayed high resilience despite receiving antibiotic treatment.

To test whether this pattern reflected broader coordination between the gut and airway microbiota, we computed Mantel correlations between airway and faecal Jaccard distance matrices at each sampling time point (Figure 7E). Correlations were consistently positive, although generally modest. The strongest association was observed immediately after antibiotic administration and pathogen challenge (D+1; *r* ≈ 0.50). Correlations subsequently declined but remained positive throughout the study (*r* ≈ 0.1–0.4). Coupled to the dysbiosis, antibiotic treatment induced marked taxonomic restructuring in both compartments, characterised by depletion of dominant resident taxa and expansion of opportunistic microorganisms in the airways, along with loss of key anaerobic fermentative and SCFA-producing bacteria in the gut (Additional Files 12 and 13).

#### Antibiotic-induced microbiota restructuring is accompanied by microbial metabolic shifts

Antibiotic-induced microbiota restructuring was accompanied by marked changes in microbial metabolic activity, with SCFAs showing the most pronounced responses (Figure 8A). Acetate was significantly affected by the antibiotic × time interaction (*p* = 0.017). Propionate was influenced by antibiotic treatment (*p* = 0.023) and showed a strong time-dependent response (*p* = 0.0019). Likewise, butyrate varied over time (*p* = 0.0004) and was strongly modulated by antibiotic exposure (*p* = 0.0010). Valerate followed a similar pattern, with significant effects of both antibiotic treatment (*p* = 0.017) and the antibiotic × time interaction (*p* = 0.027). In contrast, branched-chain SCFAs were comparatively stable, showing only trend-level associations for isobutyrate (*p* ≈ 0.075) and isovalerate (*p* ≈ 0.085).

**Figure 8.**
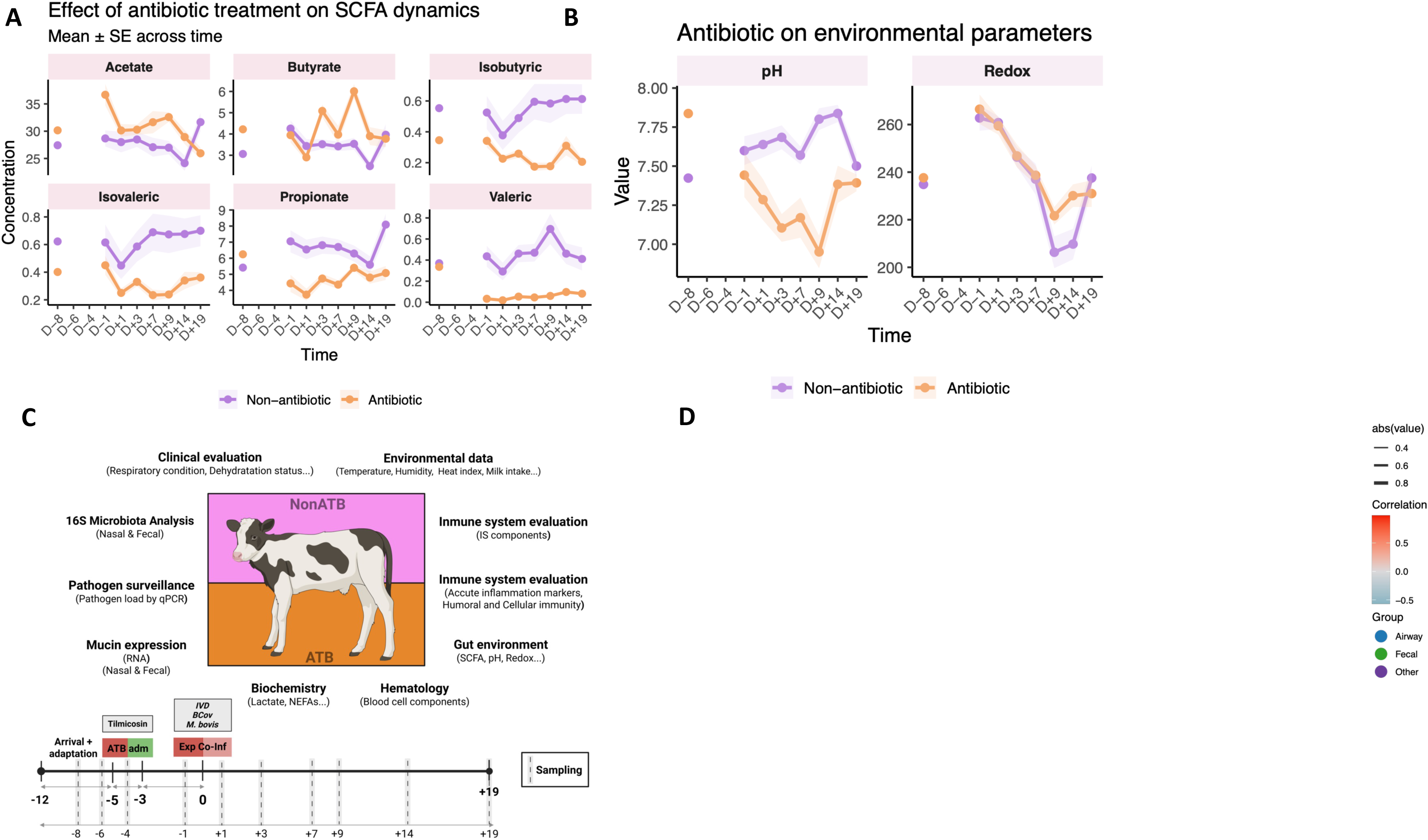
Integrated experimental design and multi-compartment analysis of the effects of antibiotic treatment on host–microbiota interactions in calves. **(A)** Effect of antibiotic treatment on short-chain fatty acid (SCFA) dynamics. Mean ± SE concentrations are shown over time for major (acetate, propionate, butyrate) and minor SCFAs; **(B)** Temporal dynamics of environmental parameters (pH and redox potential). Mean ± SE values are shown for antibiotic-treated and untreated groups; **(C)** Overview of the experimental design and sampling strategy. Calves were assigned to either a non-antibiotic (NonATB) or antibiotic-treated (ATB) group receiving tilmicosin. Samples and clinical measurements were collected before, during, and after experimental coinfection challenge. The study integrated clinical evaluation, environmental measurements, microbiota profiling, pathogen quantification, mucin expression analysis, biochemistry, haematology, and immune-system assessments from nasal and faecal compartments, and **(D)** Correlation network integrating airway mucins (blue nodes), faecal mucins (green nodes), SCFAs an environmental parameters (purple nodes), and immune, haematological, and other variables (dark purple nodes). Edges represent significant correlations, with red and blue lines indicating positive and negative associations, respectively. Edge thickness is proportional to correlation strength.

Consistent with SCFA analyses, PERMANOVA of KEGG orthologue profiles demonstrated that antibiotic exposure was the primary driver of the putative functional variation in both airway (R² = 0.276, *p* = 0.001) and faecal microbiomes (R² = 0.065, *p* = 0.001). The strongest functional shifts involved pathways associated with microbial carbon utilisation and energy metabolism. Across both ecosystems, antibiotic treatment altered glycolysis, pyruvate metabolism, the tricarboxylic acid (TCA) cycle, oxidative phosphorylation, and several fermentation-related pathways, including propanoate and butanoate metabolism (Additional Files 14 and 15). Beyond central metabolism, antibiotics consistently affected pathways involved in microbial interactions with the mucosal environment. Among the most strongly modified pathways were amino sugar and nucleotide sugar metabolism, ABC transporters, quorum sensing, bacterial secretion systems, biofilm formation, and two-component regulatory systems. Although mucin production is host-derived, many of these pathways contribute to mucin glycan utilisation, bacterial adhesion, biofilm establishment, and adaptation to mucus-associated ecological niches. Notably, the parallel responses observed in both the respiratory and intestinal microbiotas suggest that antibiotic exposure modifies common ecological processes that govern microbial colonisation of mucosal surfaces across the gut-airway axis. Consistent with the strong selective pressure imposed by antimicrobial exposure, several antibiotic resistance-associated pathways were enriched among the most responsive functional categories, including β-lactam resistance in the respiratory microbiota and monobactam biosynthesis and CAMP resistance in the intestinal microbiota.

Lastly, changes in microbial metabolism were accompanied by modifications of the intestinal physicochemical environment (Figure 8B). Faecal pH varied over time (*p* = 0.0024), decreased with antibiotic treatment (*p* = 0.0116), and showed a strong antibiotic × time interaction (*p* = 1.7 × 10□□). In contrast to SCFAs, pH was also associated with pathogen load (*p* = 0.016). Redox potential exhibited a different pattern, being driven almost exclusively by temporal dynamics (*p*= < 2 × 10 ¹□□), with no detectable effects of antibiotic treatment (*p* = 0.31), pathogen abundance (*p* = 0.63), or their interaction (*p* = 0.23; Figure 8B).

### Multi-omics integration identifies a three-stage gut-lung response to antibiotic and pathogen perturbation

We then integrated clinical, immunological, haematological, mucosal, metabolic, and microbial datasets using correlation analysis, network inference, ENVFIT, and DIABLO (Figure 8C).

Correlation analyses identified two principal modules (Additional Figure 1). The first module comprised airway and intestinal mucins, forming highly cohesive, compartment-specific clusters characterised by strong intra-compartment correlations (typically *r* = 0.6–0.8). Consistent with the limited cross-compartment mucin associations observed previously, correlations between respiratory and intestinal mucins remained weak (*r* = 0.1–0.3). The second module was centred on immune, inflammatory, metabolic, and ecological variables and was characterised by a denser network of cross-system interactions. Within this second module, SCFAs emerged as the principal connector variables linking host and microbial responses. Acetate, propionate, butyrate, and valerate showed positive associations with microbial diversity in both airway and intestinal communities (*r* ≈ 0.3–0.5) and correlated with immune markers and mucin expression (*r* ≈ 0.2– 0.4). In contrast, pathogen burden occupied a different position within the network, showing strong positive relationships with inflammatory markers (*r* ≈ 0.3–0.6) and negative associations with microbial diversity (*r* ≈ −0.3 to −0.5). Network topology further reinforced this organisation, with immune markers and SCFAs occupying the most-connected positions, while mucin variables formed peripheral yet internally cohesive subnetworks (Figure 9D).

Complementary, temporal ENVFIT analyses revealed that these interactions followed a highly structured temporal succession (Figure 9A–B). Prior to infection (D−8), microbial communities were moderately associated with mucosal and inflammatory variables (R² = 0.20–0.40, *p* < 0.05), reflecting the baseline organisation of the gut and airway ecosystem. Immediately after antibiotic administration (D−1), microbiota composition became increasingly associated with respiratory tract mucins. Airway communities were strongly linked to MUC1, MUC4, MUC16, and MUC20 (R² = 0.35–0.55, *p* < 0.01), whereas faecal communities were associated with MUC5AC, SAA, and immune-cell populations (R² = 0.30–0.50, *p* < 0.05).

Following infection (D+1 to D+7), immune and inflammatory variables became the dominant drivers of microbiota organisation. In the airways, community composition was significantly associated with pathogen burden, SAA, γδ T cells, CD4 cells, IgA, and IgM (R² = 0.30–0.60, *p* < 0.05). Faecal communities showed even stronger relationships with inflammatory and immune variables, including TNF-α, SAA, and γδ T cells (R² up to 0.60, *p* < 0.01).

From D+9 onwards, the strongest explanatory variables shifted from immune markers to microbial metabolites. In the intestinal ecosystem, acetate, propionate, butyrate, and valerate collectively explained 30-60% of community variation (R² = 0.30-0.60, *p* < 0.01), surpassing most inflammatory variables. Concurrently, airway communities progressively reverted towards a mucin-associated organisation.

DIABLO integration independently supported these conclusions. The factors that best discriminated antibiotic-treated from untreated animals were not microbial taxa alone, but coordinated combinations of microbial, immunological, mucosal, and metabolic variables (Figures 9CD). In the respiratory compartment, the most discriminative signature combined SAA, monocytes, neutrophils, leukocyte counts, adaptive immune markers, airway mucins (particularly MUC1, MUC4, MUC16, MUC20, MUC21, and MUC5AC), and SCFAs. These variables were tightly associated with respiratory taxa implicated in airway ecology and disease, including *Moraxellaceae* members. In the intestinal compartment, the dominant signature centred on SCFAs, inflammatory proteins, IgM, IgG, IgA, γδ T cells, CD45, CD4, and intestinal mucins, particularly MUC12 and MUC16, along with major fermentative genera such as *Prevotella*, *Megamonas*, *Megasphaera*, *Acidaminococcus*, *Sutterella, Biophila*, and *Intestinimonas*.

## DISCUSSION

In the present study, we combined longitudinal measurements of pathogen abundance, mucin expression, immune and physiological responses, microbial ecology, microbial metabolism, and gut environmental parameters to characterise holobiont dynamics during experimental bovine respiratory disease, with particular emphasis on the ecological consequences of antibiotic treatment, a cornerstone of BRD control.

Our findings reveal three major principles. First, respiratory pathogen load primarily drove clinical disease and acute inflammation rather than large-scale microbiota disruption in the airways and digestive tract. Second, antibiotic exposure emerged as the dominant ecological perturbation affecting microbial communities, microbial metabolic function and mucin expression. Third, microbial fermentation products, particularly SCFAs, emerged as central integrative variables linking microbiota dynamics, mucosal regulation, and immune responses across the gut–airway axis. These observations collectively support a sequential model of antibiotic-induced ecological destabilisation, infection-driven immune activation, and SCFA-associated recovery. More broadly, they suggest that while antibiotics remain essential tools for controlling BRD pathogens, their benefits should be weighed against potential ecological costs, including disruption of microbial ecosystem stability and altered mucosal homeostasis.

One of the most consistent findings was that pathogen abundance strongly tracked clinical severity and acute inflammatory responses. Serum amyloid A emerged as the host variable most closely associated with respiratory pathogen load, whereas adaptive immune markers displayed substantially weaker relationships. SAA and haptoglobin are recognised as the major acute-phase proteins in cattle [54] and are among the most sensitive biomarkers of respiratory disease and systemic inflammation [55]. Numerous studies have shown that SAA levels rise rapidly during BRD and often correlate more closely with disease progression than conventional clinical indicators [54, 56, 57]. Interestingly, SAA in our study was associated with total pathogen load rather than with any individual infectious agent. This observation reinforces the concept that acute-phase responses integrate cumulative inflammatory pressure rather than pathogen-specific signals. In multifactorial diseases such as BRD [42], where clinical severity frequently arises from interactions among viruses, bacteria, and host responses, integrated inflammatory markers may therefore provide a more biologically relevant representation of disease burden than individual pathogen measurements [58]. In contrast, most adaptive immune responses were influenced primarily by temporal trajectories rather than pathogen-load gradients. Leukocyte infiltration (CD45 cells), and CD4 T cells exhibited marked post-infection dynamics, whereas only NK cells and humoral markers (IgA, IgG and IgM) showed a significant association with pathogen abundance. These findings suggest that innate inflammatory mechanisms remain closely linked to the current infectious status, whereas adaptive immune responses are driven predominantly by longer-term immune programming and maturation, although antibody levels may still reflect antigenic load [59]. Such temporal dissociation between acute inflammatory and adaptive immune responses has been repeatedly observed during respiratory viral infections [54, 60, 61] and is considered a hallmark of the successful immune transition from pathogen control to tissue recovery.

One of the most unexpected findings was the relatively limited ecological impact of pathogen burden compared with antibiotic exposure. However, this observation should be interpreted cautiously, as natural transmission of the experimental pathogens reduced the separation between inoculated and non-inoculated calves, potentially attenuating infection-associated effects. Despite this limitation, antibiotic treatment consistently explained a larger proportion of the variation observed in microbial diversity, community composition, biomass, ecological drift, community stability, and resistance. This finding aligns with recent ecological models of microbiome perturbation, which view antibiotics as ecosystem engineers that can restructure microbial communities independently of disease status [62, 63]. Antibiotic exposure can reduce diversity, eliminate keystone taxa, alter metabolic networks, and create ecological opportunities for opportunistic microorganisms [64]. Importantly, these effects often persist after pathogen clearance, producing long-lasting alterations in community assembly and ecosystem resilience [65].

Our data demonstrate that the antibiotic effects were not restricted to the gut. Oral administration of tilmicosin, a macrolide antimicrobial with preferential distribution to bovine respiratory tissues and activity against major BRD pathogens [66, 67], induced coordinated changes in both respiratory and intestinal microbiota, supporting the concept that microbial perturbations can propagate across multiple mucosal ecosystems simultaneously. This observation matters because most microbiome studies remain compartment-focused, despite growing evidence that mucosal communities are connected through systemic immune and metabolic pathways [68]. While some calves remained relatively stable despite infection and antimicrobial exposure, others exhibited marked perturbations in both the respiratory and intestinal microbiota. The simultaneous instability in respiratory and intestinal ecosystems suggests that resilience may reflect a systems-level property of the host–microbiome assemblage rather than a characteristic of a single microbial niche [69–71]. This interpretation is consistent with growing evidence that recovery from diet- or antibiotic-induced perturbation is highly individualised [72, 73] and influenced by host factors, baseline microbiota composition, and ecological interactions.

A particularly novel aspect of this study is the characterisation of airway and faecal mucin biology during BRD and antibiotic treatment. Although mucus is widely recognised as a critical component of respiratory and gut defence [6, 8], detailed investigations of mucin dynamics in cattle remain extremely limited. We demonstrated that airway and intestinal mucosal barriers are organised according to fundamentally different architectural principles. The respiratory compartment was dominated by membrane-associated mucins, particularly MUC1, MUC4, MUC16, MUC20, and MUC21, whereas the intestinal system was overwhelmingly structured around secreted MUC2. This observation is fully consistent with current understanding of mucin biology. Hoorens et al. [17]identified nine membrane-associated mucins in cattle, including MUC1, MUC4, MUC16, MUC20, and MUC21, and showed that membrane-bound mucins are a major component of mucosal epithelial surfaces, where they contribute to barrier integrity, pathogen sensing, and host-microbe interactions. In contrast, secreted gel-forming mucins, particularly MUC2, are characteristic components of the gastrointestinal mucus layer and are fundamental to physically segregating the intestinal microbiota from the epithelial surface [5, 9, 74]. The predominance of membrane-associated mucins in the respiratory tract and of MUC2 in the intestine therefore reflects the distinct functional requirements of these mucosal environments, with the lung relying primarily on epithelial surface protection and signalling, whereas the intestine depends on a thick secreted mucus barrier to maintain microbial homeostasis. The remarkable stability of MUC2 expression despite respiratory pathogen challenge and antibiotic exposure is noteworthy. While previous studies in cattle have shown that gastrointestinal nematode infections trigger extensive mucus remodelling, including increased MUC2 expression and activation of mucin biosynthetic pathways, supporting a protective role of mucus hypersecretion during parasite infection [75, 76], no such response was observed here. Despite clear evidence of intestinal microbiota dysbiosis, MUC2 abundance remained unchanged. This finding suggests that alterations in microbial community structure alone may be insufficient to drive mucus remodelling and that substantial changes in the mucus barrier likely require direct host-pathogen interactions accompanied by local inflammatory responses. This finding could reflect strong selective pressure to preserve intestinal barrier integrity despite shifts in microbiota composition and gut metabolic activity. Importantly, this does not contradict reports that antibiotics impair mucus barrier function [77]. Vancomycin has been shown to rapidly reduce gut goblet-cell mucus secretion through direct effects on host cells, independent of the microbiota, leading to mucus barrier defects. Thus, antibiotic-induced dysfunction may occur through altered mucus secretion, turnover, or organisation rather than changes in MUC2 expression itself [77]. In contrast, airway mucins exhibited extensive expression shifts in response to both antibiotics and pathogen burden. These modifications align with the established role of airway mucins in pathogen sensing, epithelial signalling, and innate immune regulation [9].

However, our results go beyond confirming known mucin expression patterns by showing that distinct mucin architectures are associated with distinct microbial ecosystems, suggesting that mucus organisation may be an important yet previously overlooked determinant of microbial community structure across mucosal surfaces. In the respiratory tract, membrane-associated mucins, particularly MUC1, MUC4, MUC16, MUC20, and MUC21, were closely associated with key airway genera, including *Moraxellaceae* genera, taxa that play central roles in bovine respiratory health and disease. Given the established functions of membrane-bound mucins in bacterial adhesion, epithelial signalling, and immune regulation, these associations suggest that mucus organisation may actively help shape respiratory microbial communities. In the intestine, MUC12 and MUC16 were associated with SCFAs and major fermentative genera, including *Prevotella*, *Megamonas*, *Megasphaera*, *Acidaminococcus*, and *Intestinimonas*. These taxa produce microbial metabolites that regulate epithelial function and immune homeostasis, highlighting a close link between mucus architecture and microbial metabolic activity. Notably, MUC16 emerged repeatedly across multiple analytical approaches and in both mucosal compartments, together with SCFAs. Although the transmembrane MUC16 has received considerable attention in epithelial biology and immune regulation [78], its role in respiratory host–microbiota interactions remains underexplored. Experimental evidence indicates that MUC16 forms a dense glycocalyx on the apical surface of epithelial cells, serves as a physical and biochemical barrier against microbial adhesion and invasion, and contributes to immune signalling and the infiltration of inflammatory cells [79], suggesting that it may act as a key interface between microbial communities and host tissues. These findings provide a valuable first framework for future mechanistic studies aimed at understanding how mucus organisation influences microbial ecology and disease resilience during BRD.

The strongest integrative signal identified throughout the study involved SCFAs. This observation’s biological importance extends beyond microbial metabolism. SCFAs are increasingly recognised as central molecular mediators of gut–lung communication [80]. Their effects include regulating epithelial barrier function, modulating immune-cell differentiation, promoting regulatory immune pathways, enhancing antimicrobial defences, and attenuating excessive inflammation [81–84]. Our findings strongly support these concepts. SCFAs consistently occupied central positions within correlation networks, were among the strongest explanatory variables in ENVFIT analyses, and represented major discriminative features in DIABLO models. Most importantly, SCFAs became the dominant explanatory variables during recovery, when inflammatory signals progressively diminished. This transition suggests that restoring microbial metabolic function is not merely a consequence of recovery but may actively contribute to re-establishing homeostasis across the gut–airway axis.

Integrating all datasets revealed a consistent temporal pattern. The initial phase was associated with antibiotic exposure and was characterised by shifts in microbiota composition, changes in mucin expression, and alterations in microbial metabolic profiles. This was followed by a phase associated with pathogen challenge, during which pathogen load, SAA, immune responses, and clinical signs became more strongly associated with ecosystem variation. During the recovery period, associations with SCFAs progressively increased, coinciding with reduced inflammatory signals and a partial restoration of microbiota-associated functions. Therefore, the temporal shift from inflammatory biomarkers to SCFAs suggests that ecosystem recovery may be driven less by pathogen elimination than by restoration of microbial metabolic capacity.

The repeated emergence of SAA, humoral markers, MUC16, and SCFAs across all analytical frameworks suggests that these variables represent a core signature of host–microbiota organisation during BRD.

### Conclusions

This study provides a comprehensive description of the calf holobiont during respiratory disease. Our results demonstrate that pathogen burden and antibiotic exposure shape the holobiont through distinct pathways: pathogen load predominantly drives acute inflammation and clinical disease, whereas antibiotics act as ecosystem-level perturbators, restructuring microbial communities, mucus barriers, and microbial metabolism. Most importantly, SCFAs emerge as pivotal mediators. Thus, these findings support the concept that resilience to bovine respiratory disease emerges from the ability of the holobiont to maintain microbial metabolic function despite infectious and antimicrobial perturbations.

### Limitations

Several limitations should be considered when interpreting these findings. First, natural transmission of *M. bovis* and BCoV resulted in partial overlap between inoculated and non-inoculated animals, although we partly addressed this by analysing pathogen burden as a continuous variable. Second, the sample size, while typical for longitudinal large-animal experiments, may have limited the ability to detect weaker associations in high-dimensional datasets. Third, mucin analyses were based on gene expression and did not assess mucin protein abundance, glycosylation patterns, or mucus biophysical properties. Fourth, 16S rRNA sequencing provided taxonomic but not direct functional information; therefore, we inferred microbial functions primarily from SCFA measurements and integrative analyses. Finally, the study design does not allow causal inference regarding the role of SCFAs or specific mucins. Future mechanistic studies are needed to determine whether these variables actively drive host– microbiota interactions during respiratory disease.

Despite these limitations, the combined longitudinal, multi-compartment, and multi-omics approach provides a comprehensive framework for understanding how infection, antibiotics, mucosal barriers, microbial ecology, and microbial metabolism interact across the bovine gut– airway axis.

## MATERIAL AND METHODS

### Ethics approval

The local animal care and use committee and the French Minister of Research reviewed and approved the study protocol for the calves’ study (reference: APAFIS #49942- 2024062000479999 v3, dated September 23rd, 2024, Ethics Committee no. 115). All protocols were conducted in accordance with the EEC Regulation (No. 2010/63/UE) governing the care and use of laboratory animals, which has been in effect in France since January 1, 2013.

### Management and housing at birth

The study included 28 Montbéliard and Holstein calves (24 males and 4 females), with a mean birth weight of 42 ± 5.1 kg. The calves were born between September and October 2024 at the Herbipole experimental farm (Macenat, France). We separated each calf from its dam immediately after birth, weighed it, and applied a unique ear tag for identification. Calves were administered approximately 1.9 L of colostrum at each of their first two feedings (within 2 hours of birth and approximately 12 hours thereafter). Within 24 hours of birth, calves were housed individually in calf hutches (Calf-Tel; Hampel Corp., Germantown, WI, and PolyDome, Litchfield, MN) bedded with wood shavings. Wood shavings were regularly replaced to maintain a dry environment. At 7.5 ± 1.3 days of age (nominally day 7), calves were transferred to wood-shaving-bedded group pens, each measuring 4.87 × 7.31 m and accommodating between eight and ten calves. Group formation was based on birth dates, with new groups initiated once each reached capacity. This process continued until all 28 calves were born.

After colostrum feeding, calves received milk replacer (Univor, Spécial Élevage, Chasseneuil du Poitou, France) prepared at 125 g/L by mixing with hot water (65°C) and fed at approximately 41°C. The milk replacer contained 63% skim milk powder, whey, partially delactosed whey, protein powder, wheat flour, brewer’s yeast extract, wheat gluten, rapeseed oil, soybean oil, and copra oil. It contained 225 g/kg crude protein (CP), 185 g/kg crude fat (CF), 3 g/kg cellulose, and 80 g/kg ash. Throughout the study, all calves were fed the same antibiotic-free calf starter. Milk replacer was administered daily in two equal meals at 0700 and 1700 h.

Bottles were cleaned and dried after each feeding. Calves were allowed to drink freely until they exhibited satiety behaviours or refused further intake. No forced feeding was implemented.

The amount of milk replacer given to each calf was precisely adjusted to meet its individual metabolic energy (ME) requirements. These requirements included the energy needed for essential functions, such as digestion, respiration, and heat production, as well as growth, represented by average body weight gain. The adaptation followed the formula:

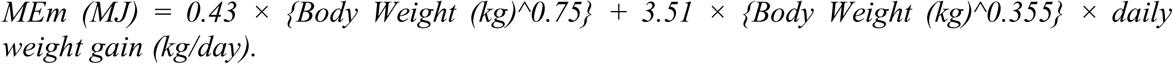

Using this formula ensured each calf received an optimal supply of metabolizable energy to support maintenance and growth throughout development. This approach achieved an average weight gain of 0.8 kg/day. Additionally, to maintain proper energy balance, we carefully regulated the environmental temperature between 15°C and 20°C.

Preventive measures included intranasal administration of the Bovine Respiratory Syncytial Virus (BRSV)- Bovine Parainfluenza-3 Virus (bPI3V) vaccine (Rispoval RS + PI3 Intranasal, Zoetis, France) at 5 days of age to safeguard against bovine respiratory disease. Additionally, halofuginone lactate (Kriptazen®, Virbac, Carros, France) was given orally at 7 days of age to manage protozoan parasites of the genus *Cryptosporidium*. Notably, the dams were not vaccinated against BCoV nor *M. bovis* to prevent the transfer of interfering antibodies during subsequent experimental infections at the ENVT facility.

### Animal handling and groups at the experimental facility

Once all 28 calves had been born at the INRAE-Herbipole experimental farm, they were transported to the research facilities of the National Veterinary School of Toulouse (ENVT, Toulouse, France). The study was conducted in a biosafety level 1 barn measuring 35 m x 11.3 m x 5 m (length x width x height) and equipped with a natural ventilation system. The barn contained four large pens (4.9 m x 14 m; total area: 73.5 m²; space per animal: 10.5 m²) in two rows. Each pen accommodated seven animals. They were all open on the front and lateral sides, with solid walls on the back and a galvanised mono-pitch roof.

At arrival (day −12), calves were 30.7 ± 13.3 days old and underwent a complete clinical examination to confirm they were clinically healthy. Animals were then balanced by body weight and sex into four experimental groups: (i) inoculated with BRDC-associated pathogens (IDV, BCoV, and *M. bovis)* without antibiotic treatment (*Inoc_NonATB*), (ii) inoculated with BRDC-associated pathogens and treated with antibiotics (*Inoc_ATB*), (iii) non-inoculated and treated with antibiotics (*NonInoc_ATB*), and (iv) non-inoculated and untreated (*NonInoc_NonATB*).

Building on this structure, we administered tilmicosin (12.5 mg/kg, twice daily; Tilmovet 250 mg/mL, Huvepharma, Antwerp, Belgium) from days −5 to −3 of the experiment. Antibiotic treatment was administered exclusively to the *nonInoc_ATB* and *Inoc_ATB* groups. Before the experimental coinoculation, all calves underwent comprehensive pathogen screening to ensure they were free of major respiratory agents. Specifically, from birth until arrival at ENVT, we monitored calves weekly for *M. bovis*, BCoV, and IDV using PCR on nasal swabs (BioT respiratory qPCR kits; BioSellal, France). We also tested them for enteric pathogens (rotavirus, Giardia*, Cryptosporidium*), with only one calf (ID18, *NonInoc_ATB*) testing positive for *Cryptosporidium*.

The two inoculated and non-inoculated pens were separated by a transparent, low-density polyethylene barrier (food-grade, 700 gsm/175 microns, density at 23 °C: 0.922 g/cm³). Within the non-inoculated or experimentally inoculated space, the two pens faced each other and were separated by a 1.5 m-wide corridor. Animals in opposing pens had visual but no tactile contact. Pens did not share equipment, food, water, space, or fomites. Each pen was treated as an independent unit, following a “*marche en avant*” system. Caretakers and students were permitted access only to either the inoculated or the non-inoculated group, but not both. Additionally, personnel and materials were never transferred between inoculated and non-inoculated groups to prevent cross-contamination. The bedding, consisting of pine shavings, was spot-cleaned daily and replaced every three days, thereby minimising dust production. All calves had *ad libitum* access to fresh water via an automatic watering system.

### Virus and bacteria preparation for the experimental co-inoculation

On day 0, calves assigned to the infected groups were co-inoculated with three BRD-associated respiratory pathogens: IDV (strain D/bovine/France/5920/2014), BCoV (strain ENVT-66), and *M. bovis* (strain RM16) [32]. The influenza D virus strain D/bovine/France/5920/2014 was isolated from the lung of a dead calf with clinical signs of BRD [85]. The virus was propagated on human rectal tumour cells (ATCC CRL-11663) in Dulbecco’s modified Eagle’s medium (DMEM; Dutscher, France) supplemented with 1 μg/mL TPCK (tosylsulfonyl phenylalanyl chloromethyl ketone)-trypsin (Thermo Fisher Scientific, MA, USA) at 37°C, 5% CO_2_ for 5 days. Viral titer was determined on swine testis cells (ATCC CRL-1746) using the 50% tissue culture infective dose (TCID_50_) method as described elsewhere [32].

The Bovine Coronavirus (BCov ENVT-66) was isolated from the lung of a dead calf with clinical signs of BRD. The virus was amplified in HRT-18G cells (CRL-3609) cultured in Dulbecco’s Modified Eagle’s Medium (DMEM; Dutscher, France) supplemented with 1 μg/mL TPCK-treated trypsin (tosyl sulfonyl phenylalanyl chloromethyl ketone; Thermo Fisher Scientific, MA, USA) at 37 °C with 5% CO for 5 days. Viral titer was determined by immunostaining on hRT18g cells using a primary monoclonal antibody (MoAb J18C-INT BCV02-12-092), followed by incubation with an Alexa Fluor 555-conjugated secondary antibody (donkey anti-mouse IgG6).

*M. bovis* strain RM16 was isolated in 2016 in France from a pool of transtracheal aspiration samples from 3 heifers with clinical signs of respiratory infection, as reported elsewhere [86]. *Mycoplasma* cells were grown in SP4 medium supplemented with ampicillin (100 μg/mL, Sigma-Aldrich) as previously described [86]. After 24-36 h of incubation at 37°C, *Mycoplasma* cultures were stored at −80°C. *Mycoplasma* titers were determined by serial dilutions in Dulbecco’s phosphate-buffered saline (DPBS; Gibco) supplemented with 2% heat-inactivated fetal calf serum (Invitrogen). Dilutions were spotted (10 μL) onto solid SP4 medium, and CFUs were counted after 2 to 5 days of incubation at 37°C. For animal inoculations, *Mycoplasma* cells were washed three times in Dulbecco’s Modified Eagle Medium (DMEM; high glucose, sodium pyruvate, and GlutaMAX-I; Gibco) by centrifugation at 9,000 *× g* for 20 min at 4 °C and kept on ice. Before inoculation, 10^10^ CFU were diluted in 10 mL DMEM. The CFUs were counted before and after the experimental infection, with no significant difference (P > 0.05).

### Inoculation of multiple pathogens via intranasal nebulization

For co-inoculation with *M. bovis*, IDV, and BCoV, we used gentle intranasal nebulization, covering the nostrils and mouth, with an adapted mask and a compressor/nebuliser system (Super Dandy Inhaler, PARI, Germany) that produces 67% of droplets below 5 µm. Qualified personnel performed nebulization with a face mask designed for foals (Swevet Piab AB, Sweden); it was non-invasive and well tolerated by the calves. Nebulization lasted 8 to 12 minutes.

The inoculation was conducted with the strain D/bovine/France/5920/2014 and the strain BCoV ENVT-66 at a dose of 10 Median Tissue Culture Infectious Dose (TCID_50_) per calf, along with *M. bovis* strain RM16 at a dose of 10¹ CFUs per calf (1 mL per animal, adjusted to a final inoculum volume of 8 mL) [32]. These doses were used in an earlier experiment [32] and successfully induced clinical respiratory signs.

### Diet

Animals were exclusively nourished with milk. The milk replacer used was the same as that provided at Herbipole (Univor, Spécial Élevage, Chasseneuil du Poitou, France), with weekly adjustments to the quantity offered to meet the energy requirements needed to achieve the target growth rate of 0.8 kg/day, as previously described. During their first two days at ENVT, calves were trained to drink milk replacer from individual buckets. Milk replacer was administered twice daily, at approximately 07:00 and 17:00. Any refusal within this time frame was recorded and quantified, and no attempts were made to force-feed calves that did not consume the provided amount. Specifically, upon arrival at ENVT, the milk replacer quantity ranged from 3 to 4.5 L per meal and gradually increased to 4.7 to 6.7 L per meal, depending on body weight. Neither concentrate nor hay was provided during the study to maintain strict control over nutrient intake and ensure that all calves received a standardised, homogeneous diet throughout the experimental period. Solid feeds introduce substantial variability in energy, fibre, and micronutrient intake, which can differ markedly between individuals depending on appetite, hierarchy, and exploratory behaviour. This variability could have introduced uncontrolled differences in physiological responses and microbiota profiles, complicating interpretation of the effects of infection and antibiotic treatment on respiratory outcomes and microbiota profiles.

### Clinical observations

The same veterinarian examined calves twice daily from arrival (Day -12) to 19 days post-infection for general condition, decreased appetite during feeding, rectal temperature, nasal discharge, coughing, abnormal breathing, respiratory rate, and abnormal lung sounds. Clinical scores were assessed for each calf as previously described with slight modifications [32]. Briefly, scores for rectal temperatures (*T*) were 0 (*T* < 39°C), 1 (39.1°C < *T* < 40°C), 2 (40.1°C < *T* < 41°C), or 3 (*T* > 41°C); scores for respiratory rates per minute (RR/min) were 0 (RR < 35), 1 (35 < RR < 40), 2 (41 < RR < 60), 3 (61 < RR < 80), or 4 (RR > 80); scores of 0 (normal), 1 (mild), or 2 (severe) were given for nasal discharge and general state; and dyspnea (absent, weak, moderate, or high) and appetite (drop in milk consumption of 0%, <30%, 30 to 60%, or > 60%) were scored from 0 to 3. We also performed thoracic ultrasonography on all calves throughout the experiment. All lung lobes were systematically examined using a standardised scanning protocol, and all animals were confirmed free of detectable pulmonary lesions before experimental coinfection.

Each calf was evaluated twice daily by two veterinarians working independently within their assigned area, providing multiple annotations per day to minimise observer bias and ensure that all labelled events corresponded to true disease-related spontaneous coughs.

### Pathogen surveillance across the study

Immediately prior to inoculation (Day 0), nasal swabs placed in a medium were analysed by real-time PCR to confirm the absence of *Mannheimia haemolytica*, *Pasteurella multocida*, *M. bovis*, *Histophilus somni*, BCoV, IDV, bovine respiratory syncytial virus (BRSV), and bovine parainfluenza virus type 3 (BPIV-3). The absence of bovine viral diarrhoea virus (BVDV) was verified by ELISA targeting the Erns antigen (IDEXX BVDV Ag/serum, IDEXX Laboratoires, Saint-Denis, France). Serological testing further confirmed that all calves were negative for *M. bovis*-specific antibodies (ELISA, Bio K 302; BioX Diagnostics) and IDV-specific antibodies (hemagglutination inhibition assay). This multi-layered screening ensured that all animals entered the study free of major respiratory pathogens, providing a clean baseline for evaluating the onset of respiratory disease under controlled infection conditions.

At days +1, +3, +7, +9, +14, and +19 following infection, animals were screened for *Mycoplasma bovis*, IDV, and BCoV using nasal swabs and bronchoalveolar lavage (BAL) fluid. Viral RNA and bacterial DNA were extracted from 170 µl aliquots of nasal swab medium, raw BAL fluid, and faecal swab samples using the NucleoMag Pathogen kit (Macherey-Nagel, Germany) on the KingFisher Flex purification system (Thermo Fisher Scientific, MA, USA) according to the manufacturer’s instructions. The PB1 (polymerase basic 1) gene of IDV, the N (nucleocapsid protein) gene of BCoV, and the oppD (oligopeptide permease D) gene of *M. bovis* were amplified using specific primers and quantified using the iTaq Universal SYBR Green One-Step kit (Bio-Rad Laboratories, USA) on a LightCycler 96 Real-Time PCR System (Roche Diagnostics, Germany). We used the bovine GAPDH gene as an internal control to assess sample quality and nucleic acid extraction efficiency. Copy numbers in samples using standard plasmids containing the PB1 product of D/bovine/France/5920/2014 (IDV), the N product of strain ENVT-66 (BCoV), or the oppD product of strain RM16 (*M. bovis*).

In parallel, at the same time points, nasal swabs were analysed using the commercial BIO-T bovine respiratory syncytial virus, parainfluenza virus type 3, *M. bovis*, *Histophilus somni*, *Mannheimia haemolytica*, and *Pasteurella multocida* PCR kits (Biosellal, France), which detect the major pathogens involved in the bovine respiratory disease complex.

### Calculation of total pathogen loads per animal

To quantify the overall infectious pressure experienced by each animal, a composite pathogen load variable was defined by integrating quantitative measurements of the three targeted respiratory pathogens: *M. bovis*, BCoV, and IDV, obtained from nasal and faecal swab samples. We expressed pathogen abundance as gene copy numbers determined by quantitative PCR assays. Given the wide dynamic range and right-skewed distribution of gene copy values, we first log -transformed all pathogen measurements after adding a pseudocount (+1) to accommodate zero values and avoid undefined logarithmic transformation. The resulting transformed variables (log [gene copies + 1]) were then summed to generate a composite metric of pathogen burden:

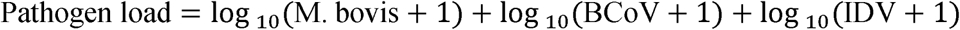

This additive approach assumes each pathogen contributes cumulatively to overall infectious pressure and integrates co-infection effects into a single continuous variable. The resulting pathogen load provides a quantitative proxy for infection intensity at the individual level.

### *Mycoplasma* cultivation from fresh swabs, DNA extraction, and whole-genome sequencing and comparative variant analysis

To determine whether the *Mycoplasma* strains detected in non-inoculated animals originated from circulation of the experimental inoculum or from an unrelated field isolate, *Mycoplasma* isolates recovered from non-inoculated animals were subjected to whole-genome sequencing and comparative variant analysis. For *Mycoplasma* isolation, nasal swabs from non-inoculated animals on days +3 and +7 were resuspended in 1 mL of SP4 medium [87], supplemented with 100 µg/mL ampicillin (Sigma-Aldrich) and 0.01% thallium acetate (Sigma-Aldrich). Samples were incubated at 37°C for 1 h and then diluted 1:10 and 1:100 in 2 mL of the same medium. After incubation at 37°C for 3 days, primary cultures were passed through a 0.22-µm filter (ClearLine) and subcultured under the same conditions. Primary cultures and subcultures were screened for mycoplasmas by PCR. Briefly, 1 mL aliquots were centrifuged at 12,000 × g for 20 min, and the resulting bacterial pellets were resuspended in 50 µL of lysis buffer containing 0.1 M Tris-HCl (pH 8.5), 0.05% Tween 20, and 0.25 mg/mL proteinase K. Samples were incubated at 37°C for 1 h, followed by proteinase K inactivation at 95°C for 10 minutes.

Samples positive for members of the genus *Mycoplasma* and specifically for *M. bovis* were identified using genus- and species-specific PCR assays, respectively [88–90].

We purified genomic DNA from the inoculum and from isolates recovered from calves 18, 23, and 25 by phenol–chloroform extraction followed by ethanol precipitation. DNA concentration and purity were assessed using a Qubit® 3.0 Fluorometer (Invitrogen, USA). Whole-genome sequencing was performed by Eurofins Genomics (Germany) using Illumina NovaSeq technology (paired-end, 2 × 150 bp).

We quality-controlled and adapter-trimmed raw sequencing reads before downstream analyses. Bioinformatic analyses were performed on the GenoToul Bioinformatics platform (Toulouse, France; http://bioinfo.genotoul.fr/). For each sample, reads were independently mapped against the *M. bovis* strain RM16 reference genome (CP077758.1) using BreSeq v0.38.1 [91] in consensus mode, with a minimum read length of 80 bp after trimming. We then compared BreSeq-generated genome difference (GD) files using gdtools COMPARE (BreSeq package) to perform comparative variant analysis and identify single nucleotide polymorphisms (SNPs) and other sequence variants shared between, or specific to, the inoculum and the recovered isolates. Whole-genome alignments and structural comparisons between the inoculum and recovered isolates were additionally performed using NUCmer (MUMmer package). Sequence alignments and identified variants were manually inspected using the Integrative Genomics Viewer (IGV v2.19.2)[92] to confirm selected variants and assess read support.

### *Mycoplasma* longitudinal cultivation from frozen swabs, DNA extraction, PCR amplification and sequencing

All nasal swabs from all animals at day −8, −1, +1, +3, +7, +9, +14 and +19 intended for *Mycoplasma* detection and isolation were stored at −80 °C prior to cultivation in standard *Mycoplasma* broth (2.45 g heart infusion broth (Difco), 90 mL Aqua bidest., 20 mL horse serum (Xebios), 10 mL fresh yeast extract, 1.2 mL Calf thymus DNA (Difco), with 20 mg thallium acetate and 1000 IU Penicillin G/mL (WDT), pH 7.8).

The nasal swabs were thawed and shaken into 2.5 mL of standard *Mycoplasma* broth, then incubated for 5 minutes at room temperature. The swabs were then pressed out and removed. A total of 0.2 mL of BAL was inoculated to 1.8 mL of standard *Mycoplasma* broth. Broth cultures were incubated at 37°C and 5% CO_2_ under static conditions and daily observation for 2 to 5 days, until turbidity was observed.

Aliquots of the broth were poured onto Petri dishes containing standard *Mycoplasma* agar. The Petri dishes were then incubated under the same conditions and checked daily for 3–10 days, until *Mycoplasma* colonies were observed. Semi-quantitative *Mycoplasma* colony counts were scored from 0 (no colonies), 1 (low grade; up to 10 colonies per plate), 2 (medium grade; 11-50 colonies per plate), to 3 (high grade; more than 50 colonies per plate).

In parallel, DNA was extracted directly from each sample without prior enrichment using the High Pure PCR Template Preparation Kit (Roche) following the manufacturer’s instructions. *M. bovis* was detected using a species-specific real-time PCR protocol [93]. Samples were categorised as positive if their Cq value was ≤36, as questionable if their Cq value was >36 and ≤38, and as negative if their Cq value was >38. Samples that were *M. bovis*-PCR negative but culture positive were retested using a pan-*Mycoplasma* PCR based on amplification of a 600 bp fragment of the 23S-rRNA gene with the PowerTrack SYBR Green Master Mix kit (Applied Biosystems) to detect other *Mycoplasma* species [94]. We identified them by Sanger sequencing the fragments (Eurofins Genomics, Ebersbach, Germany) and comparing sequences using BLAST.

To identify *Mycoplasma* spp. after enrichment, DNA was extracted from positive broths that did not yield a positive *M. bovis* PCR result by direct amplification using the High Pure PCR Template Preparation Kit. Subsequently, 23S-rRNA-based pan-*Mycoplasma* PCR, Sanger sequencing, and BLAST analysis enabled species assignment of the enriched *Mycoplasma* cultures.

### Detection of *Mycoplasma bovis*-specific antibodies

For all animals at days −8, −1, +1, +3, +7, +9, +14, and +19, serum samples were tested for *M. bovis*-specific antibodies using the ID Screen® *M. bovis* Indirect ELISA kit (Innovative Diagnostics), following the manufacturer’s instructions.

### Haematology

Blood samples were collected longitudinally from all calves at days −8, −1, +1, +3, +7, +9, +14, and +19 relative to pathogen challenge. At each time point, blood was collected into a 5 mL K_3_- EDTA vacuum tube (Venoject, Terumo, Europe NV, Leuven, Belgium) via a 0.8 × 40 mm needle (Venoject, Terumo, Europe NV, Leuven, Belgium) from the jugular vein. Immediately after sample collection, the tubes were gently mixed by 10 inversions and placed on a mechanical mixer (Spec Mix, CT6478, Drew, Oxford, USA). Blood samples were processed within an estimated timeframe of ≤2 hours after collection.

A complete automated blood count was performed on each specimen using a Sysmex XN-1000V analyser (Sysmex, Kobe, Japan). This included the following: white blood cell count (WBC); red blood cell count (RBC); haemoglobin concentration (HGB); haematocrit (HCT); mean corpuscular volume (MCV); mean corpuscular haemoglobin (MCH); mean corpuscular haemoglobin concentration (MCHC); red blood cell distribution width as coefficient of variation (RDW-CV); standard deviation (RDW-SD); reticulocyte percentage (RET%); reticulocyte count (RET); platelet count (PLT); plateletcrit (PCT); and mean platelet volume (MPV).

A blood film was prepared from each specimen, air-dried, and stained with a modified May– Grünwald–Giemsa stain (Aerospray Pro Haematology, Elitechgroup Inc., Logan, USA) before being coverslip-mounted. The smears were examined by a board-certified clinical pathologist using light microscopy (Eclipse Ci microscope, Nikon, Japan) to determine the differential leukocyte count, including the percentage of neutrophils (NEUT%), lymphocytes (LYMPH%), monocytes (MONO%), eosinophils (EO%) and basophils (BASO%).

### Blood absolute lymphocyte counts

At each time point before and after infection, blood samples were collected in 3K-EDTA tubes to determine the absolute counts of several lymphocyte subsets. Whole blood was stained using BD Trucount™ Tubes (BD Biosciences, San Jose, CA, USA) according to the manufacturer’s instructions and subsequently analysed by flow cytometry. Briefly, mix 200 µL of EDTA-anticoagulated blood thoroughly with 40 µL Transfix® (Sanbio), then dilute 1:50 in PBA (PBS containing 0.5% bovine serum albumin and 0.1% sodium azide). Next, add 20 µL of antibody cocktail to a BD TruCount tube, then add 50 µL of diluted blood. Samples were incubated for 15 min in the dark at room temperature. The antibody cocktail consisted of mouse anti-bovine CD45-AF647 (clone CC1, IgG1), mouse anti-bovine CD335-PE (clone AKS1, IgG1), mouse anti-bovine CD4-AF700 (clone CC8, IgG2a), mouse anti-bovine CD8α-PB (clone CC63, IgG2a), and mouse anti-bovine WC1-FITC (clone CC15, IgG2a), targeting leukocytes, natural killer (NK) cells, CD4+ T helper cells, CD8+ cytotoxic T cells, and γδ T cells, respectively. All antibodies were obtained from Bio-Rad. Erythrocytes were lysed by adding 450 µL of lysis buffer (BD Biosciences). Samples were acquired on a CytoFLEX LX flow cytometer (Beckman Coulter), and 8,000 counting beads were recorded per sample. Data were analysed using FlowJo v10.10.0 (Tree Star Inc., Ashland, OR, USA), and absolute cell counts were calculated based on the TruCount bead method.

### Measurement of blood Cytokines, Immunoglobulins, and acute-phase proteins

The concentrations of cytokines (IL-1β, TNF-α, IFN-γ, and IL-4) and immunoglobulins (IgG, IgM, and IgA) were determined from all calves at days −8, −1, +1, +3, +7, +9, +14, and +19. At each time point, measurements were performed using separate commercial enzyme-linked immunosorbent assay (ELISA) kits (Cloud-Clone Corp., USA). Acute phase proteins, including serum amyloid A (SAA) and haptoglobin (HP), were measured using a commercial ELISA kit (SAA) and a colourimetric assay (HP), both supplied by Tridelta Development Ltd. (Ireland).

All assays were performed strictly according to the manufacturers’ protocols. Optical densities were measured using an automated microplate reader (ELx800 Microplate Reader, BioTek Instruments, Inc., USA) with KC Junior software (BioTek Instruments).

Because some samples had insufficient serum volume, we could not analyse all samples for all parameters. The number of missing samples for each measurement was as follows: IL-1β (n = 7), TNF-α (n = 26), IFN-γ (n = 4), IL-4 (n = 100), IgG (n = 4), IgM (n = 6), IgA (n = 6), SAA (n = 6), and HP (n = 6).

### Total Antioxidant Status (TAS) and lactate

Total antioxidant status was determined in all animals and time points by LABÉO (Saint-Contest/Caen, France), an accredited veterinary and biomedical analytical laboratory. Blood samples were collected into dry tubes, and serum was separated according to standard laboratory procedures. TAS was quantified in serum using a validated automated spectrophotometric assay based on the overall antioxidant capacity of biological fluids. Results were expressed in accordance with the laboratory’s standardised methodology.

Blood lactate concentration was measured following experimental infection to assess alterations in tissue perfusion and metabolic status. Venous blood samples were collected into lithium-heparin tubes and analysed immediately after collection using a validated automated biochemical analyser at LABÉO (Saint-Contest/Caen, France). Lactate concentrations were determined using an enzymatic colourimetric method based on the oxidation of lactate to pyruvate by lactate oxidase, following the manufacturer’s recommendations and routine laboratory procedures.

### Longitudinal determination of mucins in nasal and rectal compartments

Nasal swabs were collected longitudinally from all calves at days −8, −1, +1, +3, +7, +9, +14, and +19 relative to pathogen challenge. Nasal swabs were collected into Total Nucleic Acid Preservation Tubes (Cat. 69200, Norgen Biotek), which contain an aqueous nucleic acid stabilisation buffer designed to promote rapid cell lysis, inhibit nuclease activity, and preserve RNA integrity at ambient temperature during transport and storage. RNA was subsequently isolated using the Norgen Biotek Saliva/Swab RNA Purification Kit, which employs spin-column chromatography to recover total RNA, including small RNA species.

Rectal swabs were collected in parallel using the Faecal Swab Collection and Preservation System (Cat. 45670-B, Norgen Biotek). In this system, the swab is transferred into a collection tube containing a liquid stool preservative that stabilises nucleic acids, inhibits microbial growth, inactivates pathogens, and maintains RNA integrity during transport and storage. Total RNA was extracted using the Stool Total RNA Purification Kit (Cat. 49500, Norgen Biotek), using 400 µL of preserved faecal material per extraction. This kit includes inhibitor-removal steps to eliminate compounds such as humic acids and enables simultaneous recovery of host and microbial RNA.

RNA concentration was quantified using a Qubit fluorometer (Thermo Fisher Scientific) with the appropriate RNA quantification assay (e.g. Qubit RNA HS Assay Kit).

Then, total RNA from nasal and rectal swabs was reverse-transcribed into complementary DNA (cDNA) using the Maxima H Minus First Strand cDNA Synthesis Kit (Thermo Fisher Scientific™), according to the manufacturer’s instructions, including a DNase treatment step to remove residual genomic DNA contamination.

Quantitative PCR (qPCR) was performed using the iTaq™ Universal SYBR® Green Supermix (Bio-Rad), which contains antibody-mediated hot-start iTaq DNA polymerase, dNTPs, MgCl2, SYBR Green I dye, and passive reference dyes (ROX and fluorescein). Reactions were carried out in a total volume of 5 µL, containing 1 µL of diluted cDNA and 0.4 µM of each forward and reverse primer. Amplifications were performed in 40 cycles consisting of denaturation at 95 °C for 15 s and annealing/extension at 60 °C for 1 min, following an initial denaturation step at 95 °C for 1 min. Specificity of amplification was verified by melt-curve analysis (95 °C for 10 s, 65 °C for 1 min, and 97 °C for 1 s).

A panel of 15 mucin genes was selected to capture the diversity of mucin expression across respiratory and gastrointestinal tissues. This panel included both membrane-bound (transmembrane) mucins (MUC1, MUC3A, MUC4, MUC12, MUC15, MUC16, MUC20, MUC21) and secreted gel-forming mucins (MUC2, MUC5AC, MUC5B, MUC6). The selected targets were chosen based on their reported expression patterns in the bovine upper respiratory tract (URT) and lower gastrointestinal tract (LGT) [17], allowing tissue-specific profiling of mucosal defence mechanisms.

Primer sequences were mapped to the *Bos taurus* reference genome using the Ensembl genome browser (ARS-UCD2.0 assembly; https://www.ensembl.org/Bos_taurus) to ensure species specificity and correct amplicon localisation within annotated gene regions. Despite *in silico* validation, three mucin targets (MUC8, MUC13, and MUC19) showed no or inconsistent amplification across samples, likely due to low or absent expression in the studied tissues, suboptimal primer efficiency, or incomplete genome annotation in these loci. These targets were therefore excluded from downstream quantitative analyses.

Relative gene expression levels were calculated using the 2^(-ΔCt) method. Housekeeping genes (RPL0 and RPL32) were used for normalisation after verification of stable expression across sample types. To assess residual gDNA contamination, a no-reverse-transcription (no-RT) control was included. This control consisted of a pooled RNA sample generated by combining aliquots from all samples on a plate and processing them in parallel without reverse transcriptase (replaced with nuclease-free water). The resulting no-RT control was subsequently used as a negative control in downstream analyses.

### Cross-compartment mucin association analysis

To investigate potential interactions between the airway and gastrointestinal mucosal barriers, we performed a cross-compartment association analysis that integrated airway and faecal mucin expression.

For each faecal mucin, linear mixed-effects models were fitted with each airway mucin (predictor) included individually, adjusting for faecal pathogen load, nasal pathogen load, and antibiotic treatment. Animal identity was included as a random effect to account for repeated measurements. When mixed models failed to converge or showed singularity, standard linear regression models were used as a fallback. Given the large number of pairwise tests, we adjusted p-values using the Benjamini–Hochberg method to control the false discovery rate.

### Comparison of Mucin Profiles between Tissue and Swab Samples

Total RNA was isolated from nasal turbinate samples (n = 28) and rectal tissue samples (n = 28). Briefly, approximately 2 × 2 × 2 mm tissue fragments were transferred into 350 µL of RA1 lysis buffer supplemented with 3.5 µL dithiothreitol (10 mM) from the NucleoSpin RNA Kit (Macherey-Nagel, Düren, Germany). Samples were mechanically homogenised in CKMix lysing tubes (Bertin Technologies, Montigny-Le-Bretonneux, France) in a bead-beating system until complete tissue disruption. RNA extraction was then performed according to the manufacturer’s instructions, including RNA adsorption onto a silica membrane, on-column DNase digestion to eliminate residual genomic DNA, and sequential washing steps. RNA was eluted in 60 µL RNase-free water and stored at −80 °C until analysis. RNA yield and purity were determined using a CLARIOstar microplate reader (BMG LABTECH, Offenburg, Germany) by measuring absorbance at 260, 280, and 230 nm, and results were expressed as RNA concentration (ng/µL), A260/A280, and A260/A230 ratios.

Mucin abundance was compared in paired nasal turbinate tissue and swab samples, as well as paired rectal tissue and faecal swab samples, to assess the extent to which swabs reflected mucin expression in the target tissue. Nasal and rectal swabs collected at the final sampling point (D+19) were compared to mucin expression measured in the nasal turbinate and rectal tissues collected at necropsy (D+20). Only matched samples from the same animal were included.

For each mucin, agreement between tissue and swab measurements was evaluated using Pearson’s correlation, Spearman’s rank correlation, and ordinary least squares (OLS) regression. Correlation coefficients (r and ρ), coefficients of determination (R²), and associated p-values were calculated, with statistical significance defined as p < 0.05.

To compare overall mucin expression patterns, principal component analysis (PCA) was performed using normalised mucin abundance data. Similarities between tissue and swab ordinations were further quantified by Procrustes analysis, which aligns the PCA configurations through rotation, translation, and scaling. The goodness of fit was assessed using the Procrustes statistic (m²), and significance was determined via permutation testing (10,000 permutations).

### Quantification of short-chain fatty acids in faecal samples

For short-chain fatty acid (SCFA) analysis, approximately 1 g of fresh faecal material of each animal and each time point was homogenised with 0.2 mL of 25% (v/v) sulfuric acid to stabilise volatile compounds and inhibit microbial activity. Samples were then stored at −20 °C until analysis.

Prior to extraction, faecal samples were thawed on ice and processed for gas chromatography. To separate the liquid fraction containing SCFAs, samples were centrifuged at 2,880 × g for 20 min at 4 °C. The resulting supernatant was carefully collected, and proteins were precipitated by mixing 1 mL of the supernatant with 200 µL of 25% (v/v) metaphosphoric acid, then centrifuged at 20,000 × g for 15 min at 4 °C.

For chromatographic analysis, combine 100 µL of the clarified supernatant with 75 µL of 4- methylvaleric acid (0.2% v/v) as an internal standard, then dilute to 900 µL with ultrapure water. An aliquot of 1 µL of this final mixture was injected into a gas chromatograph (Hewlett Packard 7890A) equipped with a flame ionisation detector and a DB-FFAP capillary column (30 m × 0.53 mm internal diameter, 1 µm film thickness; Agilent Technologies).

Chromatographic data were processed using Chromeleon software (Thermo Fisher Scientific). Concentrations of six SCFAs—acetic acid (C2:0), propionic acid (C3:0), butyric acid (C4:0), valeric acid (C5:0), iso-butyric acid (iso-C4:0), and isovaleric acid (iso-C5:0)—were quantified using calibration curves and internal standard normalisation. Total SCFA concentration was calculated as the sum of all measured compounds, and relative proportions were expressed as the percentage contribution of each individual SCFA to the total.

To account for variability in faecal water content, SCFA concentrations were normalised to faecal dry matter. Briefly, an aliquot of each faecal sample (5 g) was weighed (fresh weight) and dried in a forced-air oven at 60 °C for 48 h until a constant weight was reached. The percentage of dry matter was calculated as the ratio of dry weight to fresh weight. Measured SCFA concentrations (expressed per gram of fresh faeces) were then corrected by dividing by the dry matter fraction, allowing expression of SCFA levels per gram of dry faecal matter.

### Faecal and nasal microbial 16S rRNA gene sequencing

#### Faecal and nasal microbial DNA extraction

Faecal and nasal samples were collected longitudinally from all calves at days −8, −1, +1, +3, +7, +9, +14, and +19 relative to pathogen challenge and processed for microbial DNA extraction following thawing on ice. Total DNA from faecal samples was isolated using the E.Z.N.A.® Stool DNA Kit (Omega Bio-Tek, Norcross, GA, USA) according to the manufacturer’s protocol. For nasal swabs, DNA extraction was performed using the ID Gene™ Mag Fast Extraction Kit and the IDEAL 96 automated extraction platform (Innovative Diagnostics, Grabel, France) following the recommended procedures.

DNA yield and purity were assessed using both a NanoDrop spectrophotometer (NanoDrop Technologies Inc., Wilmington, DE, USA) and a Qubit 2.0 fluorometer with the dsDNA High Sensitivity assay kit (Thermo Fisher Scientific). To monitor potential contamination, negative extraction controls were included alongside all sample batches. Additional precautions were implemented throughout the workflow, including the use of UV-treated consumables, DNA-free reagents, and ultrapure water, as well as strict physical separation of pre- and post-PCR activities. We also included the ZymoBIOMICS™ Microbial Community Standard bacterial cells (Catalogue No. D6300, Zymo Research Corp., Irvine, CA, United States) (“Zymobiomics- Cells”) as a positive control during DNA extraction. According to the manufacturer’s specifications, Zymobiomics-Cells consist of three gram-negative bacteria within the phylum Pseudomonadota [*Pseudomonas aeruginosa* (theoretical composition in terms of 16S rRNA gene abundance: 4.2%), *Escherichia coli* (10.1%), and *Salmonella enterica* (10.4%)] and five gram-positive bacteria within the phylum Bacillota [*Lactobacillus fermentum* (18.4%), *Enterococcus faecalis* (9.9%), *Staphylococcus aureus* (15.5%), *Listeria monocytogenes* (14.1%), and *Bacillus subtilis* (17.4%)]. All samples were diluted to 10 ng/µL and sent to the sequencing facility.

#### Faecal and nasal 16S rRNA gene microbial sequencing

The V3-V4 hypervariable region of the 16S rRNA gene was amplified, as previously reported by our team [96–100]. Libraries were prepared according to the Illumina 16S Metagenomic Sequencing Library Preparation protocol (#15044223 RevB; Illumina Inc., San Diego, CA, USA). To minimise the risk of cross-contamination and pipetting errors, all library preparation steps were automated using a Freedom EVO NGS workstation (TECAN). Briefly, the V3-V4 hypervariable region was amplified from 10 ng of genomic DNA using Illumina-tailed primers and 2× KAPA HiFi HotStart ReadyMix (Kapa Biosystems). PCR conditions consisted of an initial denaturation at 95 °C for 3 min, followed by 25 cycles of 95 °C for 30 s, 55 °C for 30 s, and 72 °C for 30 s, with a final extension at 72 °C for 5 min. Amplicons were purified using AMPure XP beads (Beckman Coulter). Dual indices and Illumina sequencing adapters were subsequently added in a second PCR using the Nextera XT Index Kit (Illumina) and 2× KAPA HiFi HotStart ReadyMix. PCR conditions consisted of an initial denaturation at 95 °C for 3 min, followed by 8 cycles of 95 °C for 30 s, 55 °C for 30 s, and 72 °C for 30 s, with a final extension at 72 °C for 5 min. Final libraries were purified using AMPure XP beads, quantified fluorometrically using the QuantiFluor One dsDNA kit (Promega), and their size distribution was verified using a 2200 TapeStation system with DNA 1000 ScreenTape (Agilent). Library pools were quantified by qPCR using the KAPA Library Quantification Kit for Illumina platforms (Kapa Biosystems). Sequencing was performed by IGA Technology Services (Udine, Italy) on an Illumina MiSeq platform using the MiSeq Reagent Kit v3 (2 × 300 bp paired-end reads).

#### 16S rRNA Gene Sequencing Data Processing and Bioinformatics Analysis

Raw paired-end sequencing reads generated by IGA Technology Services were first subjected to quality assessment using FASTQC (v0.12.1), and the resulting reports were compiled and summarised using MultiQC (v1.27.1) to provide an overview of sequencing quality across all samples. This initial step enabled the identification of potential issues related to read quality, sequencing depth, and adapter contamination.

Primer and adapter sequences were subsequently removed using Cutadapt, as implemented in the QIIME 2 amplicon distribution (version 2024.10). After trimming, we processed high-quality reads with the DADA2 denoise-paired algorithm in QIIME 2, which performs quality filtering, error correction, chimaera removal, and amplicon sequence variant (ASV) inference to enable high-resolution reconstruction of microbial communities.

Because sequence quality and amplicon characteristics differ between sample types, we applied distinct parameter settings to faecal and nasal samples. For faecal samples, no trimming was applied to the 5′ ends of reads (trim-left = 0), and reads were truncated at 270 bp for forward reads and 250 bp for reverse reads. The maximum expected error threshold was set to 4, and a minimum overlap of 20 bp was required to merge paired reads. The error model was trained using 1,000,000 reads. For nasal (airway) samples, reads were truncated at 240 bp for both forward and reverse reads, with no trimming at the 5′ ends. The maximum expected error was set to 3.0 for both forward and reverse reads, and a minimum overlap of 10 bp was required for successful merging of paired reads.

Taxonomic assignment of the resulting ASVs was performed using a naïve Bayes classifier trained on the SILVA ribosomal RNA gene database (version 138-99-nb), allowing classification of microbial sequences across multiple taxonomic levels.

To improve data robustness and minimise the influence of low-abundance or spurious features, several filtering steps were applied. ASVs occurring only once or with very low abundance (two or fewer reads) were removed using prevalence and frequency thresholds (minimum samples = 1; minimum frequency = 3). In addition, sequences identified as mitochondrial or eukaryotic in origin were excluded from the dataset to retain only prokaryotic microbial signals.

The resulting feature table and taxonomic assignments were then imported into R (version 4.3.1; RStudio version 2025.09.1) for further analysis. Sequencing accuracy and pipeline performance were evaluated using mock microbial community controls incorporated into the sequencing. Potential contaminants were identified and removed using the decontam package (v1.28.0), based on their prevalence in negative control samples.

Following decontamination, downstream microbial community analyses were conducted in R using several packages, including phyloseq (v1.46.0) for data integration and visualisation, vegan (v2.6.4) for ecological and diversity analyses, and microbiome (v1.23.1) for compositional and structural analyses. Additional R packages were used where necessary to support statistical modelling, longitudinal analyses, and graphical representation of the data.

### Longitudinal Analysis of Microbiota Diversity and Dynamics Using Absolute Abundance Data

#### Taxonomic Aggregation and Sequence Data Summarisation

Raw Amplicon Sequence Variant (ASV) data were processed using the *phyloseq* (v1.54.0) package. A custom taxonomic fallback script was created at the genus level. ASVs with missing (NA) classification or “Unclassified” genus assignment were renamed using “Unclassified” + family-level taxonomy.

Taxonomic agglomeration was performed at the genus level. Sample depth variation was accounted for by calculating the relative abundance of each genus, normalising counts to a 0-1 scale per sample. Global metrics were calculated, including total reads, mean sequence depth, standard deviation, and total ASV identifications. Based on the ATB_Status variable, animals were split into antibiotic administration (ATB/no_ATB) groups.

#### Alpha diversity analysis

Shannon diversity and observed richness were calculated using the *estimate_richness()* function in phyloseq. Longitudinal trends in alpha diversity were evaluated across time points and experimental groups (e.g., antibiotic vs no antibiotic) and corrected for pathogen loads.

#### Beta diversity analysis

Bray–Curtis (abundance-based) and Jaccard (presence/absence-based) dissimilarity matrices were computed using the *vegdist()* function in the vegan package. Ordination of beta diversity patterns was performed using non-metric multidimensional scaling (NMDS) via the *metaMDS()* function, enabling visualisation of microbiota structure across time, treatment groups, and compartments. Ellipses representing group dispersion were added to highlight clustering patterns. Differences in microbial community composition were evaluated using permutational multivariate analysis of variance (PERMANOVA) implemented in the *adonis2()* function with 999 permutations. Models included compartment (airway vs faecal), sampling time, antibiotic exposure, and total pathogen load as explanatory variables. Statistical significance was determined based on permutation-based p-values, and the proportion of variance explained by each factor was reported as R².

To assess homogeneity of dispersion between groups, multivariate dispersion was evaluated using the *betadisper()* function followed by permutation testing (*permutest()*), as differences in dispersion may influence PERMANOVA results.

#### Beta diversity analysis with individual cohort trajectories

Temporal trajectories of microbiota structure were modelled using Principal Coordinate Analysis (PCoA) based on Bray-Curtis dissimilarities.

Longitudinal analyses were performed across all time points. Individual trajectories were visualised by connecting consecutive sampling time points with vector paths. To facilitate comparison of temporal changes, the baseline (D−8), immediate post-antibiotic exposure (D−1), and final (D+19) time points were highlighted. Cohort-level temporal shifts were assessed by calculating centroids as the mean ordination coordinates of all animals at each time point. Overlaying weighted centroid trajectories onto individual trajectories within faceted treatment panels enabled direct comparison of inter-individual variability with overall cohort dynamics throughout the study period.

#### Microbiota stability over time

For each individual, dissimilarity between successive time points and/or relative to a reference baseline (D−8) was computed using Jaccard or Bray–Curtis distances. This approach enabled estimation of microbiota stability, with low distances indicating temporal stability and high distances indicating ecological disruption. Stability metrics were summarised across time and groups to identify periods of perturbation and recovery.

#### Cumulative drift from baseline

To further quantify temporal dynamics, microbial divergence from baseline was calculated as cumulative dissimilarity relative to the earliest time point (D−8) for each individual. For each subject, a reference baseline sample was identified, and Jaccard dissimilarity between the baseline and subsequent time points was computed.

These baseline-referenced distances were analysed using linear mixed-effects models with the same structure as described above: ‘

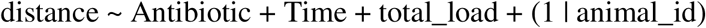

Separate models were fitted for airway and faecal compartments.

Longitudinal effects of antibiotic exposure, time, and pathogen load on microbial drift and stability were evaluated using linear mixed-effects models implemented in the lme4 package, with individual identity included as a random intercept to account for repeated measures, and statistical significance assessed using type III analysis of variance with Satterthwaite’s approximation as implemented in the lmerTest package.

#### Nasal and Faecal Microbiota co-variation

Microbiota concordance between airway and faecal compartments was assessed in R using ecological analyses implemented in the *vegan* package. Community dissimilarities were calculated independently for each compartment using Bray-Curtis and Jaccard distance metrics. Global concordance between airway and faecal microbiota structures was evaluated using Mantel tests (*mantel*, vegan), based on Spearman’s rank correlations with 999 permutations. Analyses were conducted both across the entire dataset and separately at each sampling time point using paired airway and faecal samples collected from the same animal.

Structural agreement between compartments was further investigated using Procrustes analyses (*procrustes*, *protest*, vegan) applied to NMDS ordinations generated with *metaMDS*, with significance determined by permutation testing.

To complement these community-level analyses, coordinated temporal dynamics were evaluated using a co-drift framework. For each animal and compartment, microbiota drift was calculated as the dissimilarity between each sample and its baseline community (D−8), thereby quantifying cumulative ecological change over time. Airway and faecal drift values were paired by animal and sampling day, and co-drift was assessed using Spearman correlations at each time point.

#### Definition of dysbiosis

Dysbiosis and resilience were defined using the distribution of individual mean drift values within each compartment. Animals with mean drift values ≥75th percentile were classified as dysbiotic, whereas those with values ≤25th percentile were classified as resilient. Remaining individuals were considered intermediate. To characterise microbiota trajectories, linear models were fitted to individual drift profiles, and slopes were used to identify recovering (negative slope), persistent (positive slope), or stable (near-zero slope) dynamics.

Cross-compartment comparisons enabled classification of animals as dysbiotic in both compartments, compartment-specific dysbiotic, or broadly resilient. Associations between antibiotic exposure and dysbiosis status were assessed using Pearson’s Chi-squared tests (*chisq.test()*, *stats* package).

To quantify the magnitude of microbiota perturbation following antibiotic treatment, a dysbiosis score was calculated separately for airway and faecal microbiota. Because antibiotics were administered between D−5 and D−3, samples collected at D−8 and D−6 were used to estimate baseline variability. For each animal, baseline variability was defined as the mean Jaccard dissimilarity during the pre-treatment period, while post-treatment drift corresponded to the mean Jaccard dissimilarity observed from D−1 to D+19. The dysbiosis score was calculated as:

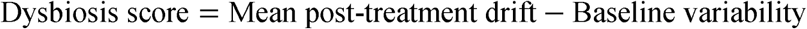

Positive values indicate divergence exceeding natural baseline variability, whereas values close to zero indicate microbiota resilience. A global dysbiosis score was obtained by summing airway and faecal scores. Animals were ranked according to this combined value, and individuals above the 75th percentile were considered the most dysbiotic.

#### MaAsLin2 Analysis of Antibiotic-Associated Microbial Signatures

To identify bacterial genera independently associated with antibiotic treatment, multivariable association analyses were performed separately for airway and faecal microbiota using MaAsLin2 (Microbiome Multivariable Association with Linear Models, version 1.16.0). Genus-level abundance tables were generated from phyloseq objects after taxonomic agglomeration at the genus rank.

For each compartment, microbial abundances were modelled as a function of antibiotic exposure while adjusting for potential confounding effects of sampling time and pathogen burden. Because samples were collected longitudinally from the same animals, we included Animal_ID as a random effect to account for repeated measurements. The following model was applied:

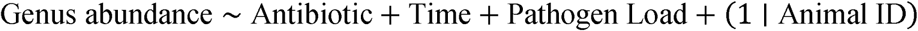

Relative abundance data were normalised using total sum scaling (TSS) and log-transformed prior to model fitting. Statistical significance was assessed using MaAsLin2’s generalised linear modelling framework, and multiple-testing correction was performed using the Benjamini– Hochberg false discovery rate (FDR) procedure. Genera with an adjusted *q*-value < 0.05 were considered significantly associated with antibiotic treatment. Positive coefficients indicate genera enriched in antibiotic-treated animals, whereas negative coefficients indicate genera depleted following antibiotic administration. The resulting models identified bacterial taxa specifically affected by antibiotic exposure, independent of temporal dynamics and infection burden, distinguishing direct antibiotic-associated microbial changes from those driven by disease progression.

#### Integrative analysis, mucus, pathogen, host response and microbiota

Integrated host–microbiota interactions were first investigated using a correlation-based systems biology approach implemented in R. The final dataset included microbial diversity indices (Shannon), mucin expression profiles, immune markers, metabolic parameters, and pathogen load. Pairwise associations were assessed using Spearman rank correlations (*cor()* with method = “spearman”, use = “pairwise.complete.obs”), which are robust to non-normal distributions and nonlinear relationships. Correlation matrices were generated using the Hmisc and corrplot packages in R, and P-values were corrected for multiple testing using the Benjamini–Hochberg false discovery rate (FDR) procedure. Significant correlations (FDR-adjusted *P* < 0.05) were retained for downstream analyses. To simplify the interaction landscape and highlight robust associations, correlations were filtered to retain only strong relationships (|Spearman ρ| > 0.35). These filtered relationships were used to construct a network representation using the igraph and ggraph packages. Nodes represented biological variables and were categorised into airway, faecal, or systemic groups based on their suffix or biological identity. Edges were defined by the strength and direction of correlations, excluding self-correlations. A force-directed layout (Fruchterman–Reingold algorithm; ggraph(layout = “fr”)) was applied to spatially organise nodes according to connectivity. Edge colour encoded correlation direction (blue for negative, red for positive; scale_edge_color_gradient2 with midpoint = 0), while edge width was proportional to absolute correlation magnitude (scale_edge_width(range = c(0.3, 2))). Node colour represented biological compartment (airway: blue, faecal: green, systemic: purple), and labels were added using repulsive text placement (geom_node_text(repel = TRUE)) to improve readability. This integrative framework enabled identification of compartment-specific interaction modules as well as cross-compartment links indicative of gut–lung axis dynamics.

Relationships between environmental, physiological, and microbial variables and overall microbial community composition were further investigated using environmental fitting (*envfit*) implemented in the vegan package. A quality filter was applied, excluding variables with >30% missing values or zero variance across all animals and time points. Residual missing values were imputed using the median, and retained numeric variables were standardised using the z-score. The ordination analysis was calculated based on the β-diversity from the microbiome. Bray-Curtis dissimilarity matrices and PCoA were reused from the β-diversity section.

The *envfit* function was applied to the PCoA vectors to project environmental variables onto the ordination space. Permutation tests (9999 permutations) were conducted to assess the statistical significance (p < 0.05) of each environmental variable. The magnitude and directional association of each physiological vector were scaled by 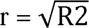.

The analysis was conducted for each time point independently across the study period. Drivers were mapped on the ordination plot as vectors. The arrow length, thickness, and direction represent the weight and orientation of the correlation with bacterial community composition. Significant variables were plotted in colours based on variable category. Non-significant variables were plotted without being coloured.

To identify coordinated patterns across multiple datasets and uncover latent structures linking host and microbial responses, Data Integration Analysis for Biomarker Discovery using Latent Components (DIABLO) was performed using the mixOmics package. Abundance microbial data were agglomerated at the genus level, transformed to relative abundances, and filtered to remove near-zero-variance features. Explanatory variables were grouped into 12 functional data blocks: Microbiome, Acute Inflammation, Haematology, Adaptive Immune Cells, Humoral Response, Cell Turnover, Metabolic Stress, Pathogen Load, SCFAs, pH, Redox, and Mucins. Prior to multi- block integration, variables with more than 30% missing values were excluded (NA > 30%).

A custom design matrix was constructed to define pairwise relationships among data blocks. Connection weights between the Microbiome block and each host physiological block were assigned according to pairwise partial least squares (PLS) correlation strengths, scaled between 0.2 and 0.4. Connections between non-microbiome blocks were assigned a baseline off-diagonal weight of 0.1.

Model hyperparameters were optimised using 5-fold cross-validation repeated three times with parallel processing (*BiocParallel*, v1.44.0). The optimal number of variables selected per component (keepX) was identified by evaluating a constrained parameter grid and maximising classification performance based on the centroid distance metric (centroids.dist). The final multi- block model was fitted using two latent components (ncomp = 2).

## Data Availability declaration

16S rRNA gene sequences from the nasal cavity and faeces are available in the DDBJ/EMBL/GenBank under BioProject ID PRJNA1517313 at the NCBI Sequence Read Archive (SRA) (www.ncbi.nlm.nih.gov/sra/PRJNA1517313). The temporary accession submission ID is SUB16425776. The associated metadata for this project are available in the INRAE institutional data repository, powered by Dataverse, and are publicly available as of the date of publication (DOI: https://doi.org/10.57745/QJFTEX). The lead contact can provide any additional information required to reanalyse the data reported in this paper upon request.

## Acknowledgments

We sincerely thank the postdoctoral researchers, technicians, and veterinary students who ensured the animals’ daily care and welfare throughout the experiment. In particular, we are grateful to Corto Martin-Rebouissou, Romain Bonsergent, Valentin Puech, Christelle Camus and Chloé Saada for their dedication and support during the study. We also thank Herbipole infrastructure for technical support.

We thank the MIGALE bioinformatics facility for providing support, computing, and storage resources (https://doi.org/10.15454/1.5572390655343293E12; Jouy-en-Josas, France).

## Author Contribution

SM assisted with statistical analyses, data integration and generated most of the figures. MM performed bioinformatic processing of the 16S rRNA sequencing data. GD performed the qPCR analyses of respiratory mucins and conducted the pilot assay to optimise primer selection. TP performed the pathogen-specific PCR assays. MC conducted the clinical examinations of the animals. DBC contributed expertise on mucin selection and primer design and advised on PCR optimisation. FZS prepared the BCoV and IDV inocula and assisted with the experimental work. AG performed the haematology analyses. GP and HR participated in DNA extraction from nasal swabs. XN performed the whole-genome sequencing analysis of the *M. bovis* isolate. CC, XN, EB, and ER provided critical feedback on the manuscript. CJ conducted cytometry and cell counts. KD performed the humoral and acute immune response analyses. CS carried out the *Mycoplasma* analyses, including culture and sequencing. EB prepared the *M. bovis* inoculum, calculated its antibiotic resistance, and isolated *M. bovis*. GM and MD supported NM in project conception, funding acquisition, and overall project direction. NM drafted the manuscript.

## TABLES

**Table 1.**
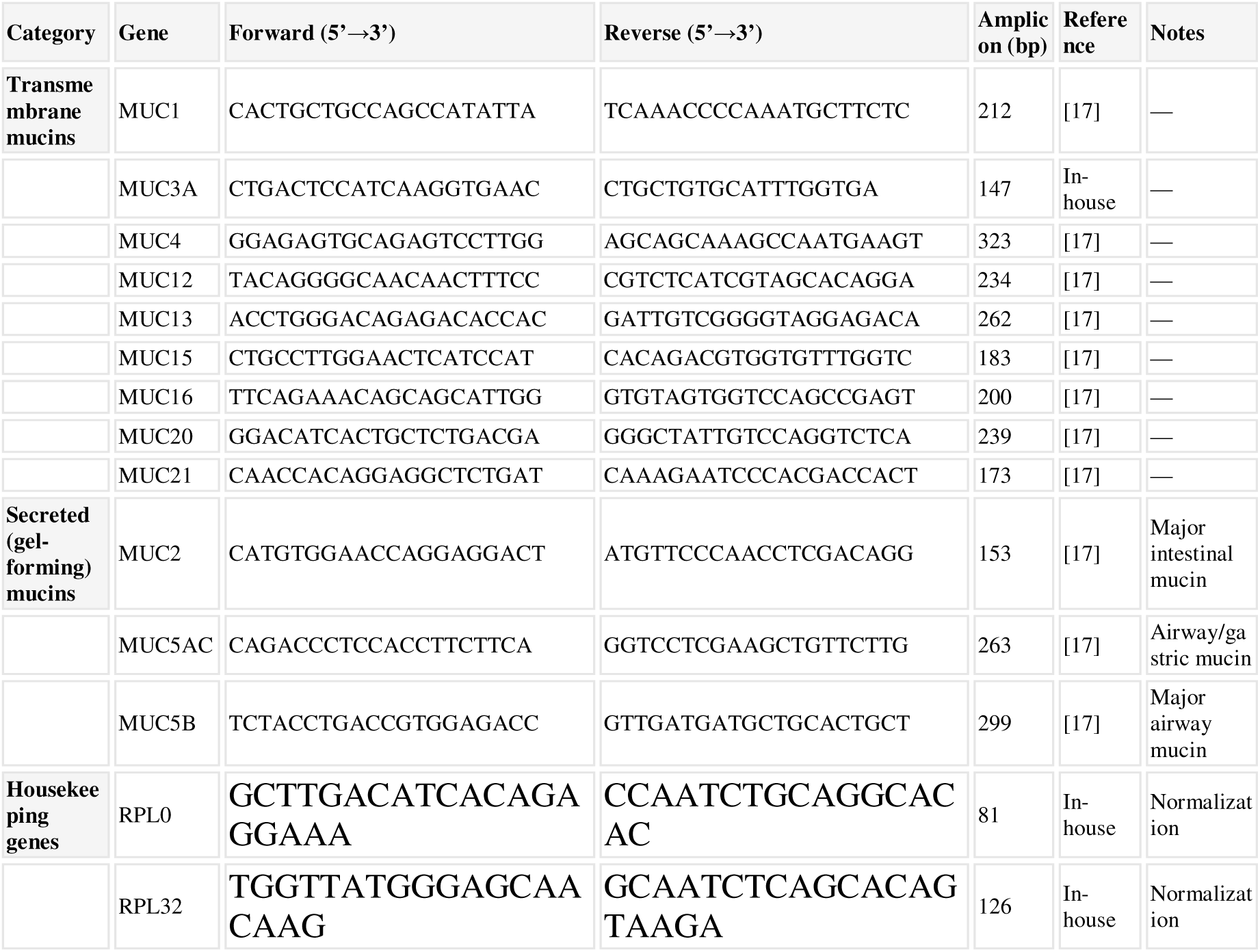
Mucins RT-qPCR primers.

## SUPPLEMENTARY FIGURES

**Additional Figure 1. Integrated microbiota-host interaction landscape across the respiratory and intestinal compartments.** Heatmap showing pairwise Spearman correlation coefficients (ρ) between mucin gene expression, microbial diversity indices, pathogen load, short-chain fatty acids (SCFAs), immune cell populations, inflammatory biomarkers, and haematological parameters measured in the airway (nasal) and faecal compartments. Correlation coefficients are represented by a blue-to-red colour scale, indicating negative and positive associations, respectively, with intensity proportional to correlation strength (Spearman’s ρ ranging from −1 to 1).

## SUPPLEMENTARY TABLES

**Additional File 1. Study metadata and measured variables.**

Comprehensive dataset containing animal demographics, management information, sampling metadata, growth performance, antibiotic exposure, clinical health scores, haematological and biochemical parameters, respiratory and faecal pathogen detection results, microbial burden measurements, short-chain fatty acid concentrations, immune cell counts, and expression levels of all mucins and immune markers included in this study.

**Additional File 2. Respiratory and enteric pathogen burden measurements.** This file contains quantitative pathogen burden data used throughout the study, including individual pathogen loads detected in nasal and faecal samples. Respiratory pathogens include *Mycoplasma bovis*, bovine coronavirus (BCoV), and influenza D virus (IDV). The dataset also includes calculated total nasal pathogen burden, total faecal pathogen burden, and overall pathogen burden (combined nasal and faecal loads), which were used to investigate associations between microbial exposure and mucin expression.

**Additional File 3. Mucin expression in the airways and faeces according to antibiotic treatment status.**

This file summarises the mean ± standard deviation (SD) of normalised mucin expression levels for all mucins analysed in this study. Values are presented in four groups: airway mucins from antibiotic-treated calves, airway mucins from untreated calves, faecal mucins from antibiotic-treated calves, and faecal mucins from untreated calves.

**Additional File 4. Airway bacterial amplicon sequence variants (ASVs).** This file contains the complete airway microbiota dataset, including all detected amplicon sequence variants (ASVs), their taxonomic assignments from phylum to species level where classification was possible, and the corresponding sequence counts for each individual animal and sample.

**Additional File 5. Faecal bacterial amplicon sequence variants (ASVs).** This file contains the complete faecal microbiota dataset, including all detected amplicon sequence variants (ASVs), their taxonomic assignments from phylum to species level where classification was possible, and the corresponding sequence counts for each individual animal and sample.

**Additional File 6. Airway bacterial genera abundance by sampling time point and antibiotic treatment.**

This file presents the mean abundance ± standard deviation (SD) of airway bacterial genera at each sampling time point, stratified by antibiotic-treated and untreated calves. For each genus, summary values are provided separately for the antibiotic and non-antibiotic groups across all study time points, enabling evaluation of temporal changes in airway microbiota composition and the potential effects of antimicrobial treatment on dominant and subdominant bacterial taxa.

**Additional File 7. Faecal bacterial genera abundance by sampling time point and antibiotic treatment.**

This file presents the mean abundance ± standard deviation (SD) of faecal bacterial genera at each sampling time point, stratified by antibiotic-treated and untreated calves. For each genus, summary values are provided separately for the antibiotic and non-antibiotic groups across all study time points, enabling assessment of temporal changes in the faecal microbiota and the potential influence of antimicrobial treatment on gut bacterial community composition.

**Additional File 8. Core airway microbiota composition.**

This file contains the genera that comprise the core airway microbiota, defined by a relative abundance threshold of ≥ 0.01% and a prevalence of ≥ 50%. For each genus, the mean relative abundance, standard deviation (SD), and prevalence (%) across airway samples are provided. Moreover, high-prevalence airway (HPA) genera and low-prevalence airway (LPA) genera, representing taxa that were broadly distributed across animals versus those detected in a smaller proportion of samples, are also described.

**Additional File 9. Core faecal microbiota composition.**

This file contains the genera that comprise the core faecal microbiota, defined by a relative abundance threshold of ≥ 0.01% and a prevalence of ≥ 50%. For each genus, the mean relative abundance, standard deviation (SD), and prevalence (%) across faecal samples are provided. We also describe high-prevalence faecal (HPF) genera and low-prevalence faecal (LPF) genera, representing taxa broadly distributed across animals versus those detected in a smaller proportion of samples.

**Additional File 10. Airway microbial alpha diversity metrics.**

This file contains alpha-diversity measurements calculated for each airway sample, including Observed ASVs (richness), Chao1 richness, Shannon diversity index, Simpson diversity index, Inverse Simpson index, and Pielou’s evenness. These metrics characterise within-sample microbial diversity and assess temporal and treatment-associated changes in the airway microbiota.

**Additional File 11. Faecal microbial alpha diversity metrics.**

This file contains alpha-diversity measurements calculated for each faecal sample, including Observed ASVs (richness), Chao1 richness, Shannon diversity index, Simpson diversity index, Inverse Simpson index, and Pielou’s evenness. These metrics characterised within-sample microbial diversity and assessed temporal and treatment-associated changes in the faecal microbiota.

**Additional File 12. MaAsLin2 results identifying airway bacterial genera differentially abundant following antibiotic treatment.** The table reports the microbial feature (genus), metadata variable tested, comparison value, regression coefficient (coef), standard error (stderr), total number of samples analysed (N), number of non-zero observations (N.not.0), nominal *P*-value (pval), and false discovery rate-adjusted *Q*-value (qval). Positive and negative coefficients indicate genera enriched or depleted, respectively, in association with antibiotic treatment.

**Additional File 13. MaAsLin2 results identifying faecal bacterial genera differentially abundant following antibiotic treatment.** The table reports the microbial feature (genus), metadata variable tested, comparison value, regression coefficient (coef), standard error (stderr), total number of samples analysed (N), number of non-zero observations (N.not.0), nominal *P*- value (pval), and false discovery rate-adjusted *Q*-value (qval). Positive and negative coefficients indicate genera enriched or depleted, respectively, in association with antibiotic treatment.

**Additional File 14. Differentially abundant microbial functional pathways (KEGG Orthology, KO) in the airways according to antibiotic exposure.** Differential abundance analysis of predicted microbial functions was performed between antibiotic-treated (ATB) and non-treated (no-ATB) animals in the airway microbiota. The table reports KEGG Orthology (KO) identifiers, effect coefficients, standard errors, nominal and FDR-adjusted p-values (q-values), enrichment direction, and mean relative abundance in each group. Positive effect coefficients indicate enrichment in ATB animals, whereas negative coefficients indicate enrichment in no-ATB animals (depleted in ATB).

**Additional File 15. Differentially abundant microbial functional pathways in faecal samples according to antibiotic exposure.** The table reports significantly altered KEGG Orthology (KO) functions identified in the faecal microbiome between antibiotic-treated (ATB) and non-treated (no_ATB) animals. For each KO entry, we report the effect coefficient, standard error, raw *P*- value, false discovery rate-adjusted *Q*-value, direction of enrichment, and mean relative abundance (%) in each group. Positive effect coefficients indicate enrichment in ATB animals, whereas negative coefficients indicate enrichment in no_ATB animals (depleted in ATB). Functional annotations are ranked by statistical significance after multiple-testing correction and provide insights into the metabolic and functional restructuring of the gut microbiome associated with antibiotic exposure.

---

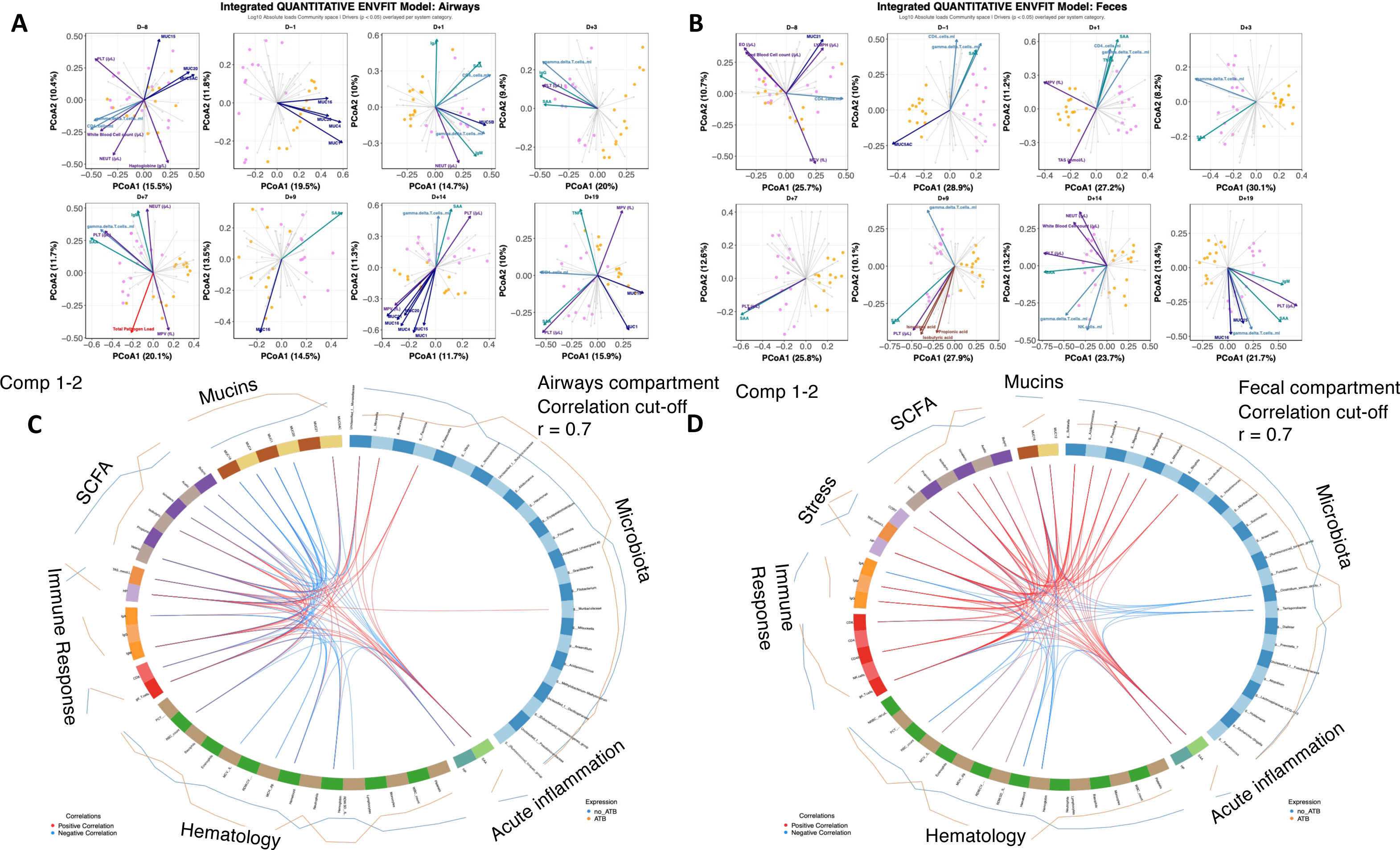

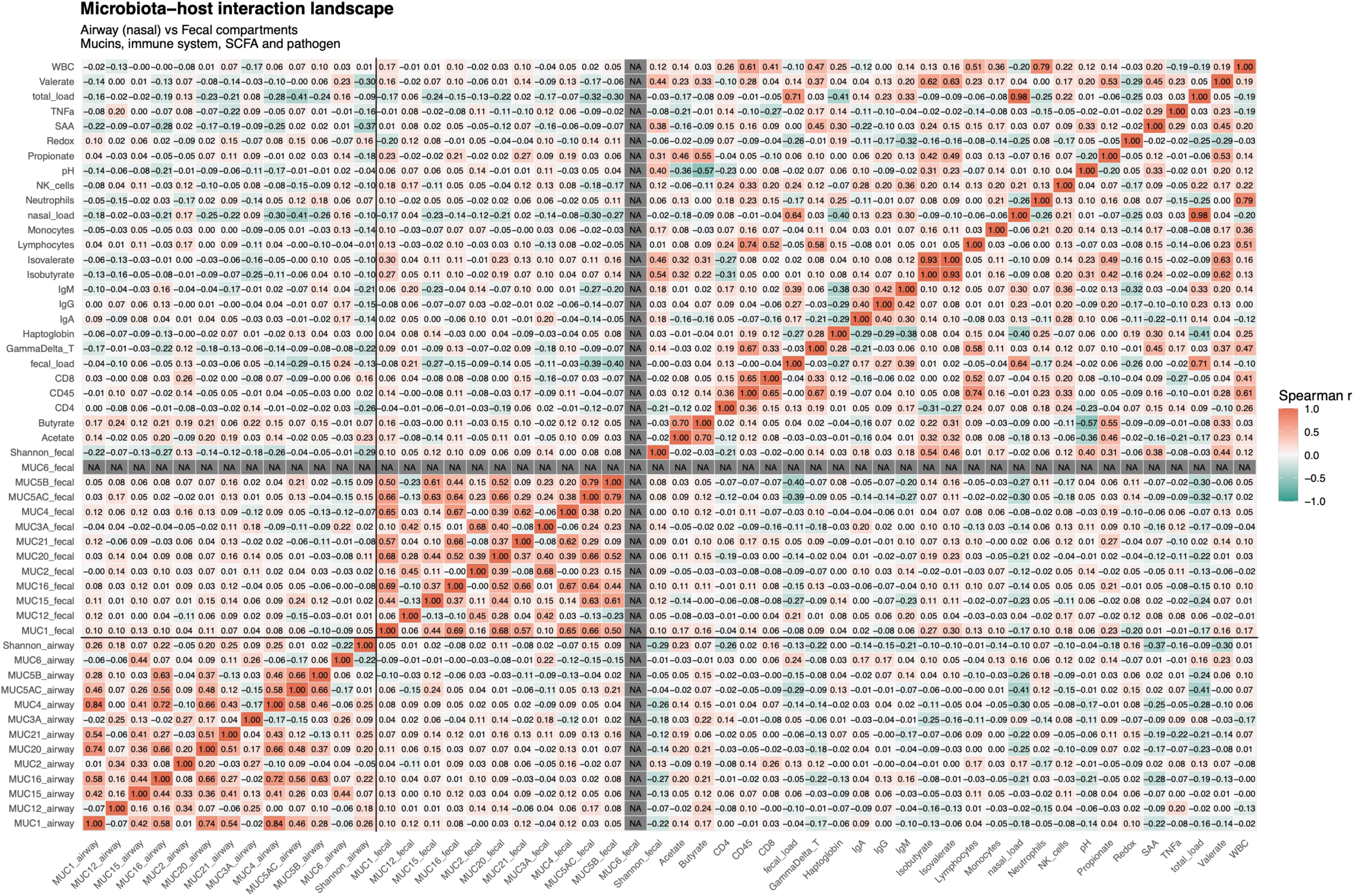

